# Trickle-down ecology: Vertical stratification of temperate forests drives asymmetric cross-layer effects on consumer communities

**DOI:** 10.64898/2026.08.04.742465

**Authors:** Jan Vigués Jorba, Manisha Bhardwaj, João Manuel Cordeiro Pereira, Anna-Lena Hendel, Daniel Kükenbrink, Lucía Villarroya-Villalba, Daniel Scherrer, Martin M. Gossner, Kurt Bollmann, Veronika Braunisch

## Abstract

1. Forests are vertically structured ecosystems where light attenuation, microclimatic buffering and resource availability occur along continuous gradients from the canopy to the understorey layers. Despite this complexity, studies rarely integrate trophic interactions across vertical layers, overlooking how vertical forest structure shapes consumer communities through abiotic and biotic pathways.
2. In this study, we combined layer-specific measurements of plant diversity, structure and microclimate with arthropod sampling in the canopy and understorey, as well as bird survey data, in a temperate forest. Applying Bayesian structural equation models with explicitly defined directional pathways, we modelled both consumer biomass and abundance across trophic levels and vertical layers.
3. Abundance measures were predominantly filtered by local layer conditions, while biomass responded to conditions across layers, reflecting stand-level energy flow. This suggests that these two metrics capture fundamentally different ecological processes. Canopy conditions consistently predicted understorey arthropod abundance across trophic levels, while the reverse was not observed, suggesting a strong asymmetric downward propagation of canopy-driven effects. Furthermore, trophic interactions between arthropod primary and secondary consumers remained largely stratified within vertical layers, suggesting a vertical food web compartmentalisation rarely shown in structurally complex aboveground systems.
4. Plant diversity, structure and microclimate shaped consumer communities mainly through the modulation of resource availability and plant apparency, with effects varying across vertical layers, trophic levels and taxonomic groups. Through complementary mechanisms, plant diversity likely determined the variability of resources available to consumers at different trophic levels. Structural properties, in contrast, potentially drove the spatial redistribution of these resources through light attenuation and microclimatic buffering, which in turn influenced the physiological capacity of consumers to access and exploit available resources.
5. These findings demonstrate that vertical stratification mediates trophic pathways in a highly directional manner, with canopy characteristics playing a disproportionate role in structuring the forest community across layers. Integrating layer-specific structural and trophic indicators into forest biodiversity assessments and management strategies is therefore essential to fully evaluate biodiversity dynamics and multifunctionality in structurally complex forest ecosystems.

## INTRODUCTION

Temperate forest ecosystems account for 17% of global forested areas (United Nations Department of Economic and Social Affairs 2026), supporting a large share of terrestrial biodiversity and providing critical ecosystem functions (Van Der Plas et al. 2018; Aerts and Honnay 2011). A defining feature of these systems is their vertical profile, where forests form a continuous gradient of light attenuation, microclimatic buffering, structural niches and resource availability (Nakamura et al. 2022; Parker 1995). Despite this inherent structural complexity, biodiversity studies often represent forest communities with metrics solely derived from understorey assessments (Bouget et al. 2011), thereby overlooking potential interactions among organisms arising from the vertical heterogeneity in niches and environmental conditions. Simplifying this complexity obscures how vegetation structure, microclimate, and biotic interactions affect forest communities across vertical layers, and how these differences propagate through organismic interactions.

Notwithstanding the increasing effort in biodiversity research to link forest structure to ecological communities, structural variables are typically aggregated across vertical layers into a single composite index (Latifi 2012). In managed European beech- (*Fagus sylvatica* L.) and Norway spruce- (*Picea abies* (L.) H. Karst.) dominated forests, the vertical profile is often functionally compressed into two distinct layers — a closed, resource-rich canopy and a deeply shaded, resource-poor understorey (Terborgh 1985). This aggregation approach can erase the within-stand gradients in ecological conditions. Furthermore, since community sampling remains mostly single-layer-based (Bouget et al. 2011), the resulting spatial mismatch between stand-level structural data and single-layer community sampling may confound the ecological significance of community interactions. Few studies assessed habitat characteristics separately by layer to test how they affect the forest community and how they mediate community interactions across temperate forest layers.

Bottom-up and top-down trophic regulation is a well-established concept in ecology. For instance, the pathway from plant community composition through arthropod assemblages to insectivorous bird populations has been extensively studied (e.g. Gruner 2004; Marquis et al. 2025). Furthermore, vertical niche partitioning within these groups is also well documented: arthropod communities differ markedly in abundance and composition between the canopy and understorey (Bouget et al. 2011; Basset et al. 2015), while bird species segregate by foraging and nesting height, especially during the breeding season (Karpińska et al. 2023).

However, vertical stratification of bird and arthropod communities have largely been studied in isolation (Šigut et al. 2018; Karpińska et al. 2023), and cross-trophic analyses integrating both groups simultaneously across vertical layers are rare (Xing et al. 2023; Sivault et al. 2024).

Over the ecological timescales relevant to animal community dynamics, resource pathways such as light interception, litter inputs, and the buffering of temperature and humidity are all processes operating predominantly asymmetrically downward through the forest profile (Ozanne et al. 2003; De Frenne et al. 2021). Integrating trophic interactions within this asymmetric physical template may reveal how canopy-driven variation in resource availability and microclimate can propagate downward through abiotic gradients, and through successive trophic levels (Vigués Jorba et al. 2026). This provides a strong mechanistic basis for expecting canopy conditions to be a disproportionate driver of community structure across the vertical profile (Yuan et al. 2012). Nonetheless, reduced resources and stable conditions in the understorey could lead to strong environmental filtering and specialisation of the plant and animal community (Carboni et al. 2016; MacArthur 1984), potentially propagating upward into canopy consumer communities through bottom-up effects of understorey plant diversity (Ebeling et al. 2018). Whether such upward effects are similar in magnitude with downward canopy forcing has, to our knowledge, not yet been explicitly tested using directional statistical frameworks.

In a system where vertical stratification not only shapes habitat structure but also energy flow and matter between trophic levels, abundance and biomass may respond differently across the vertical profile (see Ellwood and Foster 2004). These two metrics capture fundamentally different aspects of forest communities; while biomass describes trophic energy transfer (Brown et al. 2004; Lindeman 1942), abundance reflects the density of interacting individuals which is tied to processes such as predation pressure and habitat filtering (Kraft et al. 2015; Holt 2002). Consequently, biomass may be more influenced by resource quality and prey body-size distributions (Brose et al. 2006; Sterner and Elser 2002), while abundance may be more responsive to habitat heterogeneity (Wildermuth et al. 2024; Tews et al. 2004), with potential for context-dependent and interacting dynamics (Marsh et al. 2022). Whether drivers of biomass and abundance differ and whether such differences vary across trophic levels and/or vertical layers remains a poorly explored question in ecology that has direct implications for interpreting community structure and energy flow in forest food webs (Saint Germain et al. 2007).

Here, we combine layer-specific measurements of plant diversity, forest structure and microclimate with data on arthropod communities sampled in the canopy and understorey of a temperate forest, alongside bird survey data. We applied Bayesian structural equation models (BSEMs) with explicitly directional pathways and modelled consumer biomass and abundance as distinct response variables across trophic levels and vertical layers (see Fig. 1 for a conceptual review of the tested relationships). We asked the following questions: 1) do consumer biomass and abundance respond to the same drivers, or do their responses reflect distinct ecological processes, 2) do these drivers operate primarily within or across vertical layers, and 3) is there a directionality (e.g. canopy–understorey) to cross-layer effects. In addition, by investigating which specific properties of plant diversity, structure and microclimate drive consumer biomass and abundance across layers, trophic levels and taxonomic groups we provide a basis for developing layer-specific indicators that account for strata-driven processes in forest biodiversity assessments.

**Figure 1.**
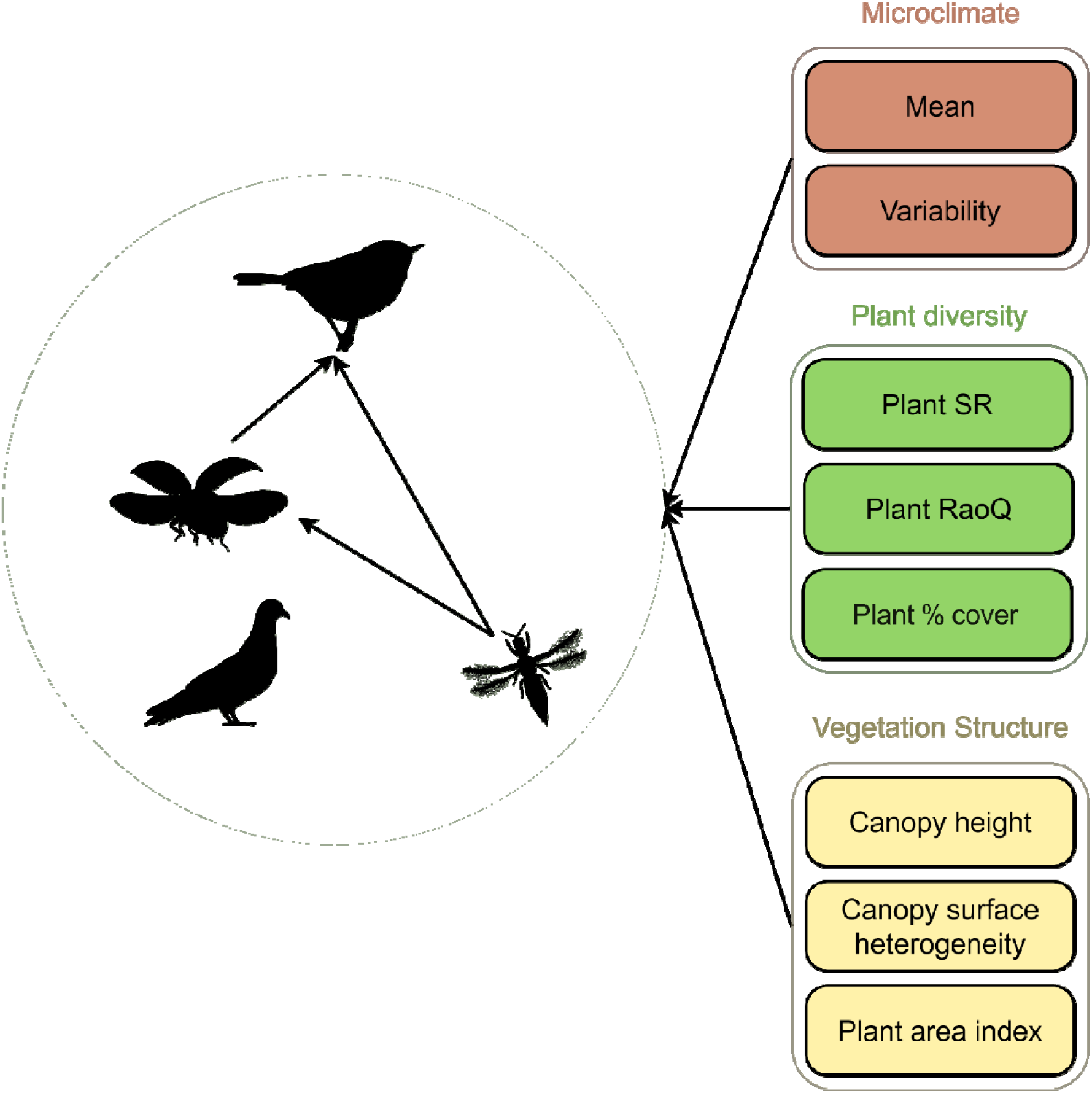
Schematic depiction of the global models used to inform the regressions included in the Bayesian SEM. Paths from each explanatory variable (here grouped into microclimatic, plant diversity and vegetation structure, right) were directly linked to each response variable within the green circle (repeated for abundance and biomass separately for the four different trophic and taxonomic groups, left). Explanatory variables of different vertical layers were allowed to coexist in the global models, prior to automated model selection and imputation into the SEM. Nevertheless, canopy height and canopy height heterogeneity were defined at the plot-level, not at the layer-level. Global models for taxonomic groups of trophic level two included the relevant response variables at trophic level one as explanatory variables. For example, this was the case for the global model of bird abundance or biomass for trophic level two (warbler silhouette, top centre), which included all explanatory variables outside the green circle as well as invertebrate (abundance or biomass respectively) at trophic level one (thrips silhouette bottom right) and two (ladybug silhouette centre left).

## MATERIALS AND METHODS

### Study site

The study took place on the Lägern mountain (47° 28′ 54″ N, 8° 22′ 37″ E) in northern Switzerland. Its highest point reaches 866 m a.s.l., with a mean annual temperature of 8.7 °C and an annual precipitation of 1102 mm (Külling et al. 2024). The area is predominantly forested and has varying land-use and management practices, with a large proportion designated as forest reserve (∼40 %), where management is either minimal or limited to measures enhancing biodiversity (swisstopo 2023). The remaining forest is managed as contiguous high forest with single or group tree selection. To cover the structural and compositional heterogeneity resulting from this, we established 33 forest plots, each covering 500 m², spanning elevations between 529 m and 820 m a.s.l. We selected plots using remote sensing data to cover the gradient of physiological (carotenoid and chlorophyll content, and equivalent water thickness) and structural (canopy height, foliage height diversity and plant area index) stand variability described in Schneider et al. (2017). The plots were located within mixed deciduous forests dominated by European beech but including other broadleaved species such as sycamore (*Acer pseudoplatanus* L.), ash (*Fraxinus excelsior* L.), and pedunculate oak (*Quercus robur* L.), while Norway spruce, silver fir (*Abies alba* Mill.) and European larch (*Larix decidua* Mill.) are the most common conifers. In each plot, a tree pertaining to the dominant stand species was designated as the focal tree (either European beech, Norway spruce or silver fir) and established as the centre of the plot.

### Field data collection

#### Plant survey

We assessed woody plant diversity once in the end of May 2024. A single expert observer identified all woody vascular plant species within the plot area and assessed their percentage cover (Dengler and Dembicz 2023) across vegetation layers. We considered two distinct layers, the understorey layer within 0.5 – 3 m and the canopy layer above 3 m (adapted from Landolt et al. 2010), while the ground layer (< 0.5 m) was excluded.

### Arthropod survey

We sampled arthropods from mid-April to early June in 2023 and 2024, using flight-interception traps (FITs), each composed of a pair of crossed transparent acrylic plates (40 cm × 60 cm) mounted above a transparent funnel leading to a collecting jar. At each plot centre, we installed one trap in the understorey (1.5 m height) and another in the canopy of the designated focal tree (mean trap height 20.6 m ± 7.6 SD). We standardised canopy measurements by sampling at the ‘centre’ of the canopy of each plot (mean tree height + mean crown base) × 0.5, following Kowalski et al. (2011), where crown base is the height at which the first leaf-bearing branch at stem rises (for further details see S1.2.2). Collected arthropods were identified to suborder-level, except for Coleoptera samples from 2023, which were identified to species-level (Table S1).

### Bird survey

Birds were surveyed by employing standardized point counts with a limited distance of 40 m from the plot centre, repeated three times per year from early April to mid-June 2023 and 2024, starting half an hour after sunrise with the latest end at 11:00 CET. Point counts consisted of a three minute adjustment period, where no birds were recorded (unless flushed away from the site), and then 10 minutes during which every bird heard or seen was recorded. As a measure of abundance, we used the maximum count of individuals per site and year.

### Microclimatic variables

To include microclimatic data in our models, at the centre of each plot we placed a temperature (EasyLog EL-21CFR-TC, Lascar Electronics, Salisbury, UK) and humidity logger (EasyLog EL-USB-2, Lascar Electronics, Salisbury, UK) in the centre of the canopy and in the understorey (∼1.5 m), set to take one measurement per hour. We calculated the hourly temperature and humidity mean and standard deviation per plot and layer across the sampling period (early April to end of June).

### Vegetation structure variables

We derived vegetation structure information from a combination of leaf-off handheld mobile (MLS) and drone-based laser scanning (ULS) acquisitions (see Table S2 for specific measures). We manually aligned the two point-cloud acquisitions and clipped this to the appropriate plot area (500 m^2^) before retrieving structural information. To obtain the mean and standard deviation of canopy height (CH and CH SD) we computed the canopy height model defined as the 99^th^ height percentile at 1 m resolution. Plant Area Index (PAI) was calculated per layer: understorey (0.5 – 3 m) and canopy (canopy base height – canopy top), as for the arthropod survey. For more details on the calculation of structural variables see S1.3.1 and S1.3.2.

### Plant diversity metrics

We calculated woody plant diversity as species richness (taxonomic diversity) and Rao’s quadratic entropy (Rao’s Q; functional diversity), for both layers separately (understorey and canopy). Rao’s Q accounts for both species’ relative abundances and functional trait dissimilarity making it particularly suited to capturing functional diversity in communities where dominant species disproportionately structure resource availability (Botta Dukát 2005). From the plant functional traits suggested by Díaz et al. (2016), we selected those influencing trophic relationships, including: leaf area (mm^2^), leaf mass per area (g m^−2^), nitrogen content per unit leaf mass (mg g^−1^) and diaspore mass (mg). All selected traits were accessed from the TRY database (Kattge et al. 2020). To compute Rao’s Q we used the R-package “FD” (Laliberté et al. 2023).

| Scale of inference | Scale at which the factor of interest is applied | Number of replicates at the appropriate scale |
| --- | --- | --- |
| Forest plot | Plot (CH and CH SD) | 33 |
| Vertical layer | Vertical layer (plant diversity, PAI and microclimate) | $33 \times 2$ |
| Arthropod community | Vertical layer | $33 \times 2$ |
| Bird community | Plot | 33 |

### Replication statement

#### Trophic level and vertical layer

To infer the contribution of each species to each trophic level, we obtained adult diet data from the open database SAviTraits (Murphy et al. 2023) for birds during the breeding period (April – June), and from Gossner et al. (2015), Chauvier-Mendes et al. (2026) and Info Fauna (2026), for arthropods. We manually imputed species-specific diet information from publicly available sources when not available from these repositories (Table S1). For omnivorous species, we kept the proportion of their adult diet belonging to each trophic level (i.e. primary or secondary consumer; decomposers were not considered). As species-level data were unavailable for non-coleopteran arthropods, we used curated regional checklists to derive a representative species sample for each suborder, from which we estimated mean trophic level proportions (Table S3, and S1.5 for further details on the checklist filtering procedure). For each suborder, we calculated the mean diet proportion at each trophic level across the filtered species pool. Data from different vertical layers were not available for birds given the difficulty of capturing this in the field, so this taxon was not analysed at the layer-level.

#### Effective abundance calculation

Abundance was taken from the count data obtained from our sampling schemes. In the case of both arthropods and birds, this was multiplied by the diet proportion at each trophic level (at the species level for birds and Coleoptera, and at the suborder level for other arthropods), to obtain a measure of effective abundance per trophic level and vertical layer. Since birds did not have layer-level data, bird effective abundance was only determined at the trophic level.

#### Effective biomass calculation

We obtained body mass data for birds from EltonTraits (Wilman et al. 2014), while mass for arthropod species was estimated across the same representative species pools described above and using allometric equations per suborder, found predominantly in Sohlström et al. (2018), but also Hawkins et al. (1997; Gastropods), Hódar (1996; Acarina and Thysanoptera), and Townsend et al. (2012; Archaeognatha) (further details on body mass sourcing in S1.7).

For non-coleopteran arthropods, body mass was estimated for each representative species within each suborder (see above), and subsequently averaged per trophic level, weighted by diet proportion (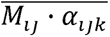, where *α_ijk_* is the species’ diet proportion at trophic level *k*, and *M_ij_* is the mass, per species *i*, in suborder *j*). This suborder- and trophic level-specific mass estimate was then multiplied by suborder abundance (*n_j_*) to yield effective biomass per suborder and trophic level (*Be_jk_*). Effective biomass values were then summed across suborders to obtain total arthropod effective biomass at each trophic level (*k*):

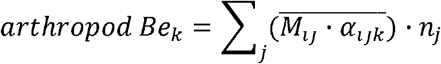

For coleopterans, biomass calculations were done at the species level (i.e. without averaging across subgroup) and added to the rest of the arthropods according to trophic level. Bird effective biomass was calculated using the same equation but at the species level, by multiplying the mass, diet proportion and abundance per species and summing the results for all species per plot and trophic level. Refer to S1.7 for further details on effective biomass calculations.

### Statistical analyses

All statistical analyses were done in the R environment (version 4.5.1, R Core Team 2022). To identify the factors driving effective biomass and abundance, we scaled all continuous variables (mean = 0, SD = 1), enabling direct comparison of coefficient estimates as measures of relative importance (Schielzeth 2010). Scaling was applied to predictor and response variables, as response variables in one regression were used as predictors in subsequent trophic levels within the BSEM. We assessed bivariate correlations among predictor variables (Table S2 and Fig. S2) and excluded variables with pairwise correlations exceeding r = 0.7.

Where multiple correlated variables were candidates for inclusion, Principal Component Analyses (PCA) loading strengths informed selection of the most representative variable. Where correlated variables were theoretically relevant and could not be reduced to a single representative, PCAs were used to integrate them into a single dimension, which was only the case for microclimatic variables (temperature and relative humidity, mean and variability) treated separately for canopy and understorey layers (S1.8, Figs. S3-S4).

Global models included predictor variables from both layers regardless of the response variable’s layer, to allow assessment of cross-layer effects (Table S4). To avoid overparameterization relative to sample size and improve model interpretability, we adopted a two-step approach in which predictor sets were further reduced using automated model selection using the ‘dredge’ function from the ‘MuMIn’ package (Bartoń 2010). Given the sample size (n = 33), predictor number was restricted to a maximum of seven to maintain a defensible ratio of observations to free parameters in a Bayesian framework and avoid overfitting (Lee and Song 2004). Model selection uncertainty was addressed by averaging coefficients across all candidate models with ΔAICc ≤ 2. Predictor variable support was quantified as the sum of Akaike weights across candidate models containing that variable (Burnham and Anderson 2002), and only variables with summed Akaike weights ≥ 0.5 were retained for subsequent structural equation modelling, as this threshold indicates a variable contributed to the majority of the cumulative model weight.

To investigate potential pathways by which structural vegetation attributes influence arthropod and bird abundance and biomass, we applied a piecewise Bayesian structural equation model (BSEM) for each response variable separately (Fig. 1), using the ‘blavaan’ package (Merkle et al. 2015), with BSEM pathways following a bottom-up direction (Fig. 1 and S1.8 for details). We fitted models using default priors on regression paths (Normal (0, 10)), with initial MCMC values drawn within this range, and used four Markov chains, each with 8000 posterior samples following a 2000 sample burn-in phase. We first evaluated model fit using modification indices (MI) after the first run, and additional paths were added when MI > 10 (Jones and Hofer 2016). We then evaluated final model fit using posterior predictive p-values (PPP), posterior predictive checks (comparing observed and simulated distributions), and R^2^ values (more details in S1.8). To further assess the robustness of our results, we repeated the statistical procedure using only Coleoptera, for which species-level identification provides the most reliable biomass estimates. Additionally, to test whether trophic links between arthropods and birds were obscured by taxa rarely consumed by birds (e.g. Acari, Thysanoptera), we repeated analyses restricting arthropods to groups known to dominate the diets of the occurring bird species (Coleoptera, Araneae, Diptera, Lepidoptera, Aphidina, Heteroptera and Holometabolic larvae). This resulted in six models, each assessing either abundance or biomass, for all arthropods, arthropods filtered by bird diet (hereafter referred to as filtered-arthropod model), or Coleoptera only.

## RESULTS

We sampled 130,089 arthropod individuals spanning 25 suborders and recorded 1,133 bird detections of 38 species, across all plots and sampling periods. Primary consumers dominated arthropod effective biomass and abundance in both the canopy and understorey, although the dominant suborders differed between the two layers and between biomass and abundance measures (e.g. Thysanoptera contributing significantly to abundance but not effective biomass; Fig. S5). Bird primary consumers also showed higher values in effective biomass than secondary consumers, but the latter were more than twice as abundant, reflecting larger body sizes of primary consumers (e.g. wood pigeon, *Columba palumbus*).

### Drivers of biomass and abundance

The BSEMs fit adequately, with PPP values across all six models ranging from 0.45 to 0.72 (Table S5) and showed strong MCMC chain convergence across all parameters. The models’ explanatory power also showed satisfactory results (mean R^2^ 0.35 ± 0.2 SD), although some individual regressions had low R^2^ values (Table S5). Below, we describe the overarching patterns across all six models (see Table S6-S8 for posterior medians and 90-95 % Equal-Tail credible intervals (CrI) across individual regressions, Fig. 3 for the full arthropod model and Figs. S6-S7 for the filtered-arthropod and Coleoptera models).

The equivalent regressions across the three model types showed largely consistent patterns across taxonomic groups. Across all models (Fig. 3, Figs. S6-S7), effective abundance of arthropods in the canopy was mostly predicted by canopy variables, while understorey abundance was predicted by both canopy and understorey variables (Fig. 2A.1). In effective biomass models arthropod canopy biomass was more frequently predicted also by understorey variables, while understorey biomass was again predicted by variables from both layers (Fig. 2A.1).

**Figure 2.**
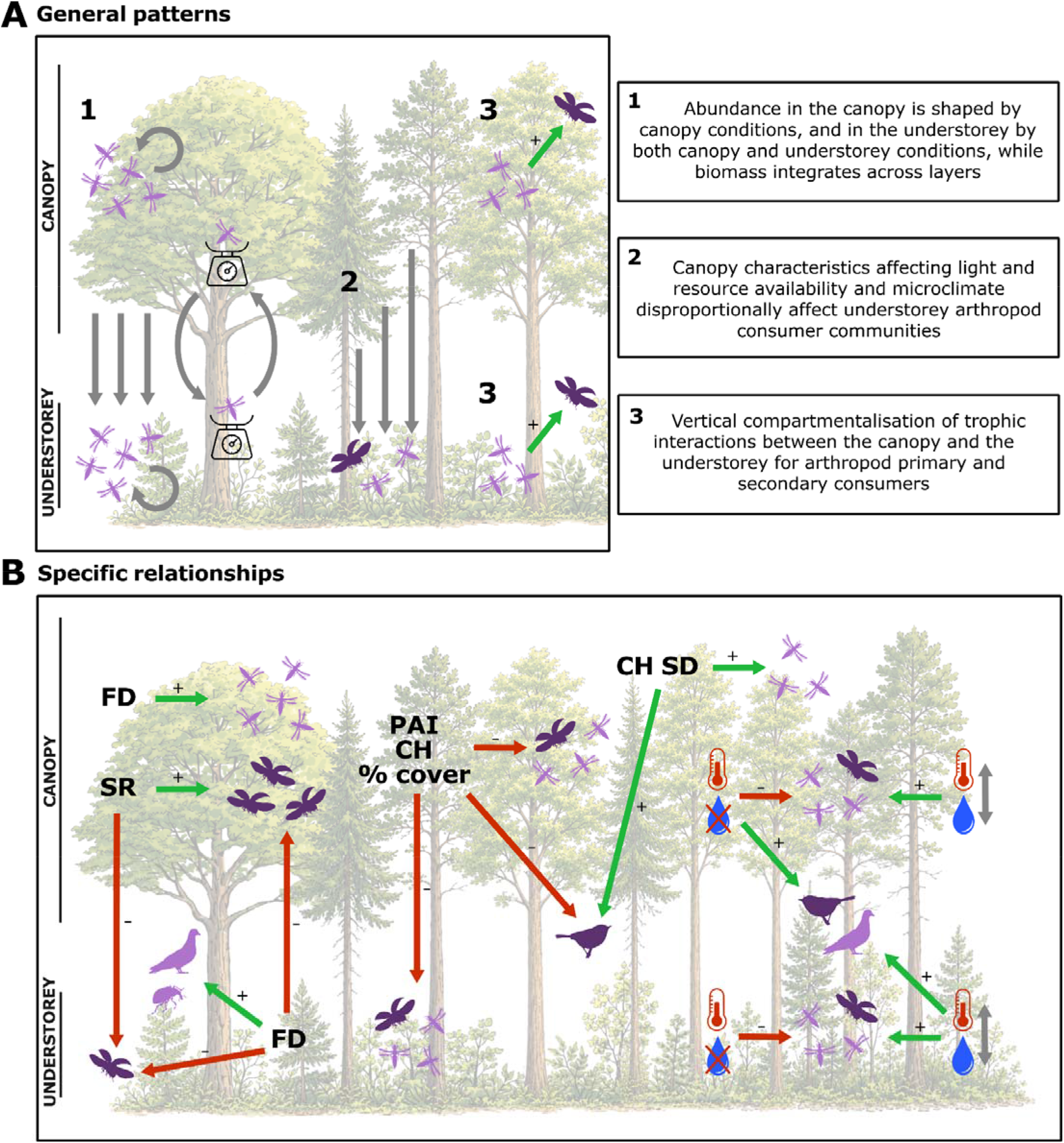
Across all panels, silhouettes represent the concerned taxonomic groups (arthropods and birds), and dark purple hues represent secondary consumers (ladybug and warbler silhouette), while light purple hues represent primary consumers (thrips, weevil (specifically representing herbivore Coleoptera) and wood pigeon). For simplicity, no distinction between biomass and abundance is made in this figure (refer to Fig. 3, S6-S7 for details). Effects are shown with arrows, where positive ones are in light green and a plus (+) sign next to the arrows, negative effects in red and a minus (−) sign, and both negative and positive in light grey and no sign. Panel A represents the generalised findings across biomass and abundance, vertical layers and trophic interactions, with numbers on the top-left corners of the boxes on the left and box line styles, linking to the text boxes on the right, with an explaining summary. This panel refers to the results addressed in the first section of the discussion. Panel B shows the detailed findings covered in the second section of the discussion, from left to right depicted in the same order as addressed in the text. Acronyms: FD (woody plant functional diversity), SR (woody plant species richness), PAI (plant area index), CH (mean plot canopy height), % cover (percentage woody plant cover), CH SD (canopy height standard deviation, i.e. heterogeneity) and symbols: red thermometer and crossed blue water droplet (mean microclimate, i.e. warm-dry axis), red thermometer and blue water droplet with up-down grey arrow (microclimate variability), are used for simplicity.

Bottom-up biotic pathways showed vertical layer consistency across most models, where abundance and biomass of arthropod primary consumers at a specific layer consistently explained that of arthropod secondary consumers at the same layer (Fig. 2A.3, Fig. 3, S6-S7). Nevertheless, the direction of the effects on effective biomass and abundance differed among vegetation-related metrics. In brief, plant Rao’s Q showed contrasting effects across trophic levels and layers, while plant species richness showed generally negative effects on arthropod biomass and abundance (Fig. 3, S6-S7). Plant cover and canopy PAI showed consistently negative effects on biomass and abundance across all arthropod trophic levels (Fig. 3, S6-S7), while understorey PAI showed mixed results, with positive effects on biomass across the arthropod biomass models and negative effects on the arthropod abundance models (Fig. 3, S6-S7), although Coleoptera models presented exceptions to this pattern (Fig. S7). Canopy height showed largely negative effects on both effective biomass and abundance across all taxonomic groups (Fig. 3, S6-S7), while canopy height heterogeneity showed consistent positive effects across all models (Fig. 3, S6-S7). Effective biomass and abundance were affected equally by microclimatic variables, with variability affecting all response variables positively across trophic levels (Fig. 3, S6-S7), while mean microclimate (high values meaning warm and dry) consistently showed negative effects on all arthropods (Fig. 3, S6-S7) and positive effects on all birds (Fig. 3, S6-S7). Refer to S2 for more detailed results and Fig. 2 for a schematic view.

**Figure 3.**
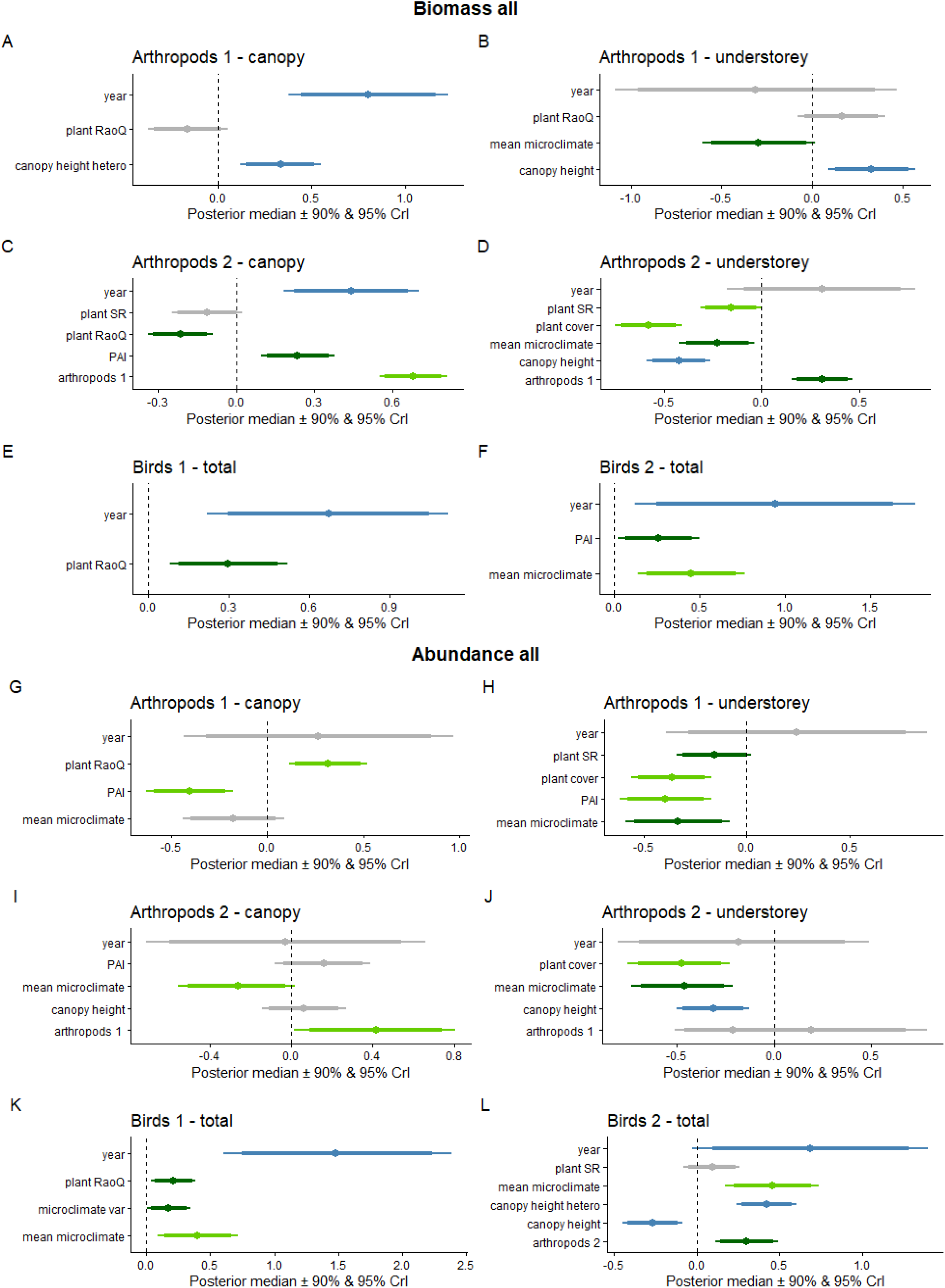
Equal-Tail credible intervals (CrIs) at 90 % (thick lines) and 95 % (thin lines, extending from thick lines), and median (central dot), for the coefficients of the variables in the BSEM. The top six panels show the predictor variable CrIs for the full biomass model and the bottom six panels those for the full abundance model. The title for each panel represents the specific regression’s predictor variable (ie. all arthropod biomass belonging to trophic level one on the top left). Coloured lines represent variables measured in the canopy (light green), understorey (dark green), or at the plot level (blue), while grey lines represent CrIs crossing zero at 90 % CrI (non-significant).

## DISCUSSION

In this study, we provide a comprehensive assessment of how vertical stratification shapes the drivers of consumer biomass and abundance across multiple trophic levels in a temperate forest ecosystem. Although our results reveal complex, context-dependent dynamics across vertical forest layers, trophic levels, and taxonomic groups, consistent patterns emerged in biomass and abundance metrics. Our study revealed three important ecological insights. First, abundance tended to be locally filtered by the corresponding layer variables, while biomass was more vertically integrated (Fig. 2A.1). This provides further evidence that biomass and abundance reflect fundamentally different ecological processes. Second, vertical stratification mediated bottom-up trophic pathways in a highly directional manner, with canopy conditions propagating asymmetrically downward to shape understorey communities (Fig. 2A.2). Third, trophic interactions between primary and secondary consumers remained largely stratified within their respective layers (Fig. 2A.3), suggesting a vertical compartmentalisation of food webs rarely demonstrated in temperate forest ecosystems.

### Vertical layers mediate effects on biomass and abundance across trophic groups

Effective abundance and biomass showed decoupled responses to the forest vertical stratification (Fig. 2A.1), which suggests that the two metrics capture fundamentally different axes of community organisation: biomass integrates the productivity and resource context at the stand-level, while abundance tracks layer-level habitat filtering. Effective biomass of arthropods showed weak stratified patterns and could be predicted by variables that were not layer specific. For instance, canopy consumer biomass was frequently predicted by understorey plant functional diversity and Plant Area Index (PAI, Fig. 3). This cross-layer signal points to biomass resulting from stand-level energy flow and productivity (Saint Germain et al. 2007). In contrast, effective abundance of arthropods in the canopy was predominantly predicted by canopy variables, while abundance in the understorey responded to both canopy and understorey predictors. In the canopy, it is likely that arthropods track local conditions for activity and resource exploitation and therefore abundance largely reflects the suitability of immediate habitat conditions rather than broader stand-scale properties (Dial et al. 2006; Wardhaugh 2014). The understorey communities, however, showed a strong canopy-driven response (Fig. 2A.2).

Canopy variables consistently predicted understorey arthropod abundance across trophic levels, while the reverse was not observed (Fig. 2A.2), indicating an asymmetric structural filtering through forest vegetation (Terborgh 1985). The downward gradient of light attenuation, plant biomass availability, and microclimate buffering through the forest profile provides the physical template for this pattern (Nakamura et al. 2022; De Frenne et al. 2021). The fact that it holds across trophic levels (i.e. not only for primary consumers) suggests that canopy-driven effects propagate through successive trophic levels. The absence of a reciprocal upward effect supports the hypothesis that in dense and strongly stratified forests the understorey does not impose equivalent constraints on canopy communities as the canopy does on the understorey, at least not regarding abundance. This pattern is consistent with the structural asymmetry inherent in systems where productivity and resource availability are concentrated in the upper stratum (Ozanne et al. 2003; Catfolis et al. 2025). This finding underscores the critical role of canopy integrity in maintaining stand-level biodiversity, reinforcing that disturbances affecting the canopy structure can have cascading impacts across trophic levels (Nakamura et al. 2017; Perry et al. 2021).

A further dimension emerging from our bottom-up pathway analysis was a consistent vertical compartmentalisation of trophic interactions between arthropod primary and secondary consumers (Fig. 2A.3). Stemming from spatial boundaries generating food web compartments (Pimm and Lawton 1980), forest vertical layers may act as distinct resource environments (Vigués Jorba et al. 2026) differentially filtering consumer communities across strata. This is particularly true for arthropods, given their limited movement capacity relative to birds, though some vertical redistribution between strata does occur (Yoshida et al. 2021). These co-occurring sub-webs along the vertical profile, as evidenced in other ecosystems such as along soil strata and water columns (Setälä and Aarnio 2002; Brewer et al. 2007), have rarely been empirically demonstrated in structurally complex aboveground systems such as forests (but see Pringle and Fox Dobbs 2008; Negrello Oliveira et al. 2025; Paniagua et al. 2009). Nevertheless, while the SEM approach suggests possible causal links, it is important to note that they rely on correlational data and lack the capacity to confirm causality. Therefore, we interpret our results as suggestive patterns and rely on further studies to improve the correlational links by studying and confirming causality.

### Drivers of effective biomass and abundance across forest layers

Plant diversity, structure and microclimate shaped abundance and biomass likely through the modulation of resource availability and plant apparency across vertical layers and taxonomic groups. Plant diversity determines the stability of resources available to consumers at different trophic levels, shaping their vertical distribution (Fig. 2B). Different complementary mechanisms may explain these patterns: the resource concentration hypothesis (Root 1973) and more-individuals hypothesis (Storch et al. 2018) predict positive diversity–abundance relationships for generalist primary consumers, while resource dilution (Muiruri et al. 2019) and the enemies hypotheses (Staab and Schuldt 2020) predict negative effects on primary consumer specialists and positive effects on secondary consumers respectively. Structural properties were influenced by the spatial redistribution of these resources through light attenuation and microclimatic buffering, in turn shaping the vertical distribution of consumers by mediating the physiological capacity of consumers to access and exploit available resources (Fig. 2B). Nevertheless, the direction of these effects was rarely consistent, varying with vertical layer, trophic level, and taxonomic group, cautioning against generalised expectations of positive diversity– and structure–productivity relationships in forest ecosystems (Forrester and Bauhus 2016; Müller et al. 2018).

Plant diversity effects were particularly complex, with contrasting outcomes across trophic levels and vertical layers. In the understorey, greater woody plant functional diversity (Rao’s Q) benefited primary consumers (particularly beetles and birds, Fig. 2B & Fig. 3, S6-7), likely by supporting a broader range of dietary resources for generalist feeders (Bernays et al. 1994; Unsicker et al. 2008), while reducing arthropod secondary consumer biomass through resource dilution decreasing prey-finding efficiency for specialists (Muiruri et al. 2019). This dilution effect appeared to extend into the canopy, where understorey plant functional diversity also negatively affected arthropod secondary consumer biomass (Fig. 2B) This suggests that bottom-up resource diversification at the base of the vegetation structure may reduce prey predictability across vertical layers (resource-concentration hypothesis, Root 1973). In contrast, higher canopy plant functional diversity may support more herbivore niches in an already resource-abundant environment, increasing primary consumer abundance following the more-individuals hypothesis (Storch et al. 2018; Wardhaugh 2014). Considering that our beech- and spruce-dominated plots maintained sufficient host plant canopy cover for specialist primary consumers (minimum 40 % beech and spruce cover; mean 66% and 63 % respectively), plant functional diversity effects likely operated through supplementing resources for generalist species (Vigués Jorba et al. 2026). The opposing effects of canopy plant species richness on secondary consumer arthropods (negative in the understorey but positive in the canopy, Fig. 2B) suggest canopy densification as the common underlying mechanism: greater plant diversity supports a more productive canopy while simultaneously reducing light penetration and understorey productivity (Dormann et al. 2020). This reflects a shift in the dominant mechanisms across layers: the enemies hypothesis explains the responses of secondary consumers in the resource-rich canopy (Staab and Schuldt 2020), while resource dilution drives the scarcity of primary consumers in the light-limited understorey (Muiruri et al. 2019).

Plant cover and PAI in the canopy consistently showed negative effects on understorey communities (Fig. 2B), reflecting reduced light availability below a dense canopy and consequently lower resource and energy availability, consistent with the metabolic theory of ecology (Brown et al. 2004; Müller et al. 2014). This negative effect was also seen in canopy response variables (Fig. 2B), likely due to flight-interception traps (FITs) placed in the centre of the canopy experiencing reduced light availability from shading of the upper canopy layers (Brüllhardt et al. 2020). These findings suggest that vertical resource gradients operate continuously rather than as discrete layer boundaries (Nilsson et al. 2022; Vinod et al. 2023).

Canopy height and heterogeneity further modulated this vertical redistribution (Fig. 2B). Taller stands concentrated resources in the canopy at the expense of the understorey, driving greater vertical stratification and negatively impacting understorey arthropod secondary consumers and Coleoptera primary consumers (Ulyshen 2011; Vigués Jorba et al. 2026). This effect may be less pronounced in younger stands where vertical layers are spatially closer.

Bird secondary consumers responding to this downward reduction in prey availability could consequently be negatively affected (Schwenk et al. 2010). Conversely, greater canopy foliage volume in taller stands likely supported higher abundances of holometabolic larvae in the canopy (in line with the “more individual hypothesis”), resulting in increased fallout into understorey FITs. This mechanism may explain the positive effect of canopy height on understorey primary consumer biomass observed during the study period (Murakami 2002).

Canopy heterogeneity had a positive effect on arthropod primary consumers in the canopy and bird secondary consumers, potentially through an increase in niche availability and resource complementarity (Loreau and Hector 2001; Storch et al. 2018).

Finally, response to microclimatic effects revealed a fundamental decoupling between arthropods and birds (Fig. 2B). On the one hand, arthropod biomass and abundance in all layers responded negatively to warm and dry conditions, contrary to established predictions (Abram et al. 2017; Brown et al. 2004). This reflects the physiological dependence of forest specialists on buffered humid environments (Enjin 2017; Pett et al. 2024) and suggests that microclimatic optima are context-dependent and specific to community-composition (Sallé et al. 2021; Decker et al. 2025). On the other hand, bird abundance and biomass responded positively to warm and dry conditions, consistent with increased foraging efficiency and activity (Maresh Nelson et al. 2024), reduced brooding energetic costs (Nord and Nilsson 2019) and lower nest-soaking rates (Wesołowski 2007). The consistently positive effect of microclimatic variability on abundance and biomass of both groups likely reflects performance optimization (Treasure and Chown 2019) and diverse microhabitat conditions supporting more functionally diverse communities through complementarity effects (Loreau and Hector 2001).

### Limitations and future considerations

Two main limitations warrant consideration. Firstly, we used subgroup biomass averages for non-coleopteran arthropods from checklist data. However, we applied extensive statistical checks and filters to mitigate biases (S1.5.1). Further, Coleoptera dominate our samples (33.7 % of effective abundance, 26.4 % of effective biomass), whereas other highly frequent groups contribute negligibly to biomass relative to their abundance (e.g. Thysanoptera (25.5 % abundance, 0.2 % biomass), see Table S3), limiting the practical impact of this caveat.

Moreover, complete local checklists were available for all remaining abundant groups except for holometabolic larvae (2.5 % abundance), which further reduced the potential bias from the use of subgroup means. Secondly, arthropod flight-interception trapping reflects activity as well as abundance, with its efficiency being subject to both climatic and structural variables (Floren et al. 2026). However, we argue that this is still appropriate, as more active individuals are also more likely to enter the trophic pathways considered in this study. The consistency of results across all six models and across both biomass and abundance metrics further strengthens the confidence in the general patterns reported here. Nevertheless, future work could refine biomass estimates with direct measurements of the sampled arthropods, and pair flight-interception trapping with complementary methods less sensitive to activity levels (e.g. direct search or branch clipping), to disentangle activity effects from true abundance and biomass patterns.

## CONCLUSIONS

Our results highlight the importance of vertical stratification as a structuring driver in temperate forest food webs. Biotic and abiotic stand characteristics differentially shaped consumer biomass and abundance through complementary ecological mechanisms and point to context-dependency rather than a generalised diversity–productivity relationship in forests. The consistent downward propagation of canopy-driven effects across trophic levels underscores the disproportionate role of canopy integrity in maintaining stand-level biodiversity. This has implications for how forest management and climate-driven canopy changes may impact multitrophic interactions and ecosystem functioning. Developing a broader spectrum of layer-specific structural and trophic indicators in forest biodiversity assessments and sustainable management strategies is therefore essential to accurately characterise biodiversity dynamics and multifunctionality in structurally complex forest ecosystems.

## Author contributions

Jan Vigués Jorba, Kurt Bollmann, Martin M. Gossner, Veronika Braunisch, João Manuel Cordeiro Pereira, Anna-Lena Hendel and Manisha Bhardwaj conceived the ideas and designed methodology; Jan Vigués Jorba, Lucía Villarroya-Villalba, Kurt Bollmann, and Daniel Kükenbrink collected and processed the data; Jan Vigués Jorba analysed the data; Jan Vigués Jorba, Kurt Bollmann, Martin M. Gossner, Daniel Scherrer, Veronika Braunisch, João Manuel Cordeiro Pereira, Anna-Lena Hendel and Manisha Bhardwaj interpreted the results; Jan Vigués Jorba led the writing of the manuscript. All authors contributed critically to the drafts and gave final approval for publication.

## Data Availability statement

Data and code used for the manuscript analyses are available on GitHub at https://github.com/jvigues/biomass-abundance-forest-layers.

## Acknowledgements

Firstly, we would like to thank Markus Gysin, Georg Rückner, Nino Tinella and Miro Keller, the tree climbers who were instrumental for setting up the canopy installations. We are also grateful to Vincent Guerra for his contribution with the vegetation surveys and Alexander Szallis for the identification of the sampled Coleoptera. We would also like to thank Mauro Marty and Peter Thür for supporting the LiDAR data acquisition.

