## Supplementary Material for "Trickle-down ecology: Vertical stratification of temperate forests drives asymmetric cross-layer effects on consumer communities"

1. METHODS
   1.1. *Study site*

The study took place on the Lägern mountain (47° 28′ 54″ N, 8° 22′ 37″ E) in northern Switzerland. Its highest point reaches 866 m a.s.l., with a mean annual temperature of 8.7 °C and an annual precipitation of 1102 mm (Külling et al. 2024). The area is predominantly forested and has varying land-use and management practices, with a large proportion designated as forest reserve (~40 %), where management is either minimal or limited to measures enhancing biodiversity (swisstopo 2023). The remaining forest is managed as contiguous high forest with single or group tree selection. To cover the structural and compositional heterogeneity resulting from this, we established 33 forest plots, each covering 500 m², spanning elevations between 529 m and 820 m a.s.l. We selected plots using remote sensing data to cover the gradient of physiological (carotenoid and chlorophyll content, and equivalent water thickness) and structural (canopy height, foliage height diversity and plant area index) stand variability described in Schneider et al. (2017). The plots were located within mixed deciduous forests dominated by European beech but including other broadleaved species such as sycamore (*Acer pseudoplatanus* L.), ash (*Fraxinus excelsior* L.), and pedunculate oak (*Quercus robur* L.), while Norway spruce, silver fir (*Abies alba* Mill.) and European larch (*Larix decidua* Mill.) are the most common conifers. In each plot, a tree pertaining to the dominant stand species was designated as the focal tree (either European beech, Norway spruce or silver fir) and established as the centre of the plot.

*1.2. Field data collection*

*1.2.1. Plant survey*

We assessed woody plant diversity once in the end of May 2024. A single expert observer identified all woody vascular plant species within the plot area and assessed their percentage cover (Dengler and Dembicz 2023) across vegetation layers. We considered two distinct layers, the understorey layer within 0.5 – 3 m and the canopy layer above 3 m (adapted from Landolt et al. 2010), while the ground layer (< 0.5 m) was excluded.

*1.2.2. Arthropod survey*

We sampled arthropods from mid-April to early June in 2023 and 2024, using flight-interception traps (FITs), each composed of a pair of crossed transparent acrylic plates (40 cm × 60 cm) mounted above a transparent funnel leading to a collecting jar. At each plot centre, we installed one trap in the understorey (1.5 m height) and another in the canopy of the designated focal tree (mean trap height 20.6 m ± 7.6 SD). To account for the heterogeneity within canopies and considering how the sampling location could affect our results, we decided to standardise canopy measurements by sampling at the ‘centre’ of the canopy of each plot, following Kowalski et al. (2011). We defined the canopy centre as (mean tree height + mean crown base) × 0.5 (Kowalski et al. 2011), where crown base is the height at which the first leaf-bearing branch at stem rises. At each site, tree climbers climbed the focal tree and set up a rope-pulley system by securing a wooden beam to the branches in the canopy with ratchet straps. We filled the collecting jars with a 0.5 % solution of Rocima^®^ GT antifungal (Lanxess AG, Cologne, NRW, Germany) and emptied and refilled them every two to three weeks. In the field, we filtered the trap contents using conventional tea filters, transferred the material into containers filled with 70 % ethanol, and stored them in a cool, dry place until laboratory processing. Arthropods were identified to suborder-level, except for Coleoptera samples from 2023, which were identified to species-level (Table S1).

*1.2.3. Bird survey*

Birds were surveyed by employing standardized point counts with a limited distance of 40 m from the plot centre, repeated three times per year from early April to mid-June 2023 and 2024, starting half an hour after sunrise with the latest end at 11:00 CET. Point counts consisted of a three minute adjustment period, where no birds were recorded (unless flushed away from the site), and then 10 minutes during which every bird heard or seen was recorded. As a measure of abundance, we used the maximum count of individuals per site and year.

*1.2.4. Microclimatic variables*

To include microclimatic data in our models, at the centre of each plot we placed a temperature (EasyLog EL-21CFR-TC, Lascar Electronics, Salisbury, UK) and humidity logger (EasyLog EL-USB-2, Lascar Electronics, Salisbury, UK) in the centre of the canopy and in the understorey (~1.5 m), set to take one measurement per hour. We calculated the hourly temperature and humidity mean and standard deviation per plot and layer across the sampling period (early April to end of June).

*1.3. Vegetation structure variables*

*1.3.1. Laser scanning acquisitions*

We derived vegetation structure information (see Table S2 for specific measures used) from a combination of leaf-off handheld mobile (MLS) and drone-based laser scanning (ULS) acquisitions. We performed MLS acquisitions using a GeoSLAM ZebHorizon (GeoSLAM Ltd, Nottingham, UK) handheld scanner following a regular grid pattern (distance between parallel lines ≈ 12.5 m) covering a 50 m x 50 m area centred around the focal tree, following Kükenbrink et al. (2025). We then processed the raw acquisitions using GeoSLAM Hub (v 6.2.1). We performed the ULS acquisitions using a Riegl miniVUX-3 scanner (Riegl, Austria) mounted on a DJI Matrice M600-pro drone, following a regular acquisition pattern consisting of two double-grids rotated by 45° flown at 60 m above ground and a distance between consecutive flight lines of 20 m (see Morsdorf et al. 2025 for more details on the ULS acquisition strategy), covering the same area as acquired by MLS. We used the Swiss CH1903+, LV95 (EPSG:2056) coordinate system and the LN02 (EPSG:5728) vertical reference system to process and georeference the raw data using the Riegl’s RiPROCESS software package (v1.9.3).

We subsequently manually aligned the two point-clouds using CloudCompare (v2.14.alpha) and clipped this to the appropriate plot area (500 m^2^) before retrieving structural information from the acquisitions. Due to restrictions on the use of drones in public areas, drone flights were not possible for two plots in deciduous stands and therefore only MLS data was used in these cases. Nevertheless, occlusion effects in the canopy from the MLS acquisitions were minimal considering that they were performed under leaf-off conditions. Furthermore, we height-normalised the point cloud to calculate canopy height, so that the height above the ground for each point was calculated using LASTool’s (v241125) *lasground* and *lasheight* functions (Isenburg 2024).

*1.3.2. Structural variable computation*

To obtain the mean and standard deviation of canopy height (the latter as a measure of canopy surface heterogeneity), we computed the canopy height model defined as the 99^th^ height percentile at 1 m resolution, based on the height-normalised point cloud (see above section) using the LASTools *lascanopy* function (v241125, Isenburg 2024). We implemented all calculations in python 3.11 in combination with the LASTools *lascanopy* function (v241125) and rasterio (v1.4.3) package. Plant Area Index (PAI) was calculated using the open-source tool AMAPVox (v2.4.1, Vincent et al. 2017). AMAPVox calculates plant area densities (PAD) at a voxel level based on the local laser attenuation by tracing all laser pulses through a predefined voxel grid. The laser attenuation per voxel was then converted to PAD using the free-path-length Maximum Likelihood Estimator described in Pimont et al. (2018). Voxel size was set to 25 cm, following Schneider et al. (2019). AMAPVox was run separately for MLS and ULS acquisitions and the resulting 3D voxel grids were combined using the AMAPVox’s *‘merge’* function. Per layer PAI was later derived from the PAD voxel grid by accumulating all PAD values for voxels above 0.5 m within a layer (understorey [0.5 – 3 m], and canopy [canopy base height – canopy top], as for the arthropod survey) and dividing by the area (500 m^2^) using AMAPVox’s *‘plantAreaIndex’* function.

*1.4. Plant diversity metrics*

We calculated woody plant diversity as species richness (taxonomic diversity) and Rao’s quadratic entropy (Rao’s Q; functional diversity), for both layers separately (understorey and canopy). Rao’s Q accounts for both species’ relative abundances and functional trait dissimilarity making it particularly suited to capturing functional diversity in communities where dominant species disproportionately structure resource availability (Botta‐Dukát 2005). From the plant functional traits suggested by Díaz et al. (2016), we selected those influencing trophic relationships, including: leaf area (mm^2^), leaf mass per area (g m^−2^), nitrogen content per unit leaf mass (mg g^−1^) and diaspore mass (mg). All selected traits were accessed from the TRY database (Kattge et al. 2020). To compute Rao’s Q we used the R-package “FD” (Laliberté et al. 2023).

*1.5. Trophic level and vertical layer*

To infer the contribution of each species to each trophic level, we obtained diet data for birds and arthropods (adult diet) from the open database SAviTraits (Murphy et al. 2023) for birds during the breeding period (April – June) and from Gossner et al. (2015), Chauvier-Mendes et al. (2026) and Info Fauna (2026), for arthropods. We manually imputed species-specific diet information from publicly available sources when not available from these repositories (Table S1). For omnivorous species, we kept the proportion of their adult diet belonging to each trophic level (i.e. primary or secondary consumer; decomposers were not considered). As species-level data were unavailable for non-coleopteran arthropods, we used curated regional checklists to derive a representative species sample for each suborder, from which we estimated mean trophic level proportions (Table S3). Suborder-specific checklists at the study region scale (canton of Zürich, records after 2000), were obtained from Info Fauna (2026) when available, and otherwise using checklists at the Swiss or Central European scale (Table S1). Checklists were subsequently filtered to retain only forest-dwelling, non-riparian, low-elevation species classified as Least Concern on the Swiss Red List (FOEN and InfoSpecies 2023), ensuring broad representativeness of the arthropod communities likely present at our study sites. All suborder species’ checklists were further filtered by body size, using length thresholds derived from our field samples: for each suborder, we identified the largest and smallest individuals captured and retained a length class as a valid threshold only if at least five individuals fell within a few millimetres of that size, ensuring the threshold reflected a common size class rather than an outlier. For each suborder, we calculated the mean diet proportion at each trophic level across the filtered species pool. Data from different vertical layers were not available for birds given the difficulty of capturing this in the field, so this taxon was not analysed at the layer-level.

*1.6. Effective abundance calculation*

Abundance was taken from the count data obtained from our sampling schemes. In the case of both arthropods and birds, this was multiplied by the diet proportion at each trophic level (at the species level for birds and Coleoptera, and at the suborder level for other arthropods), to obtain a measure of effective abundance per trophic level and vertical layer. Since birds did not have layer-level data, bird effective abundance was only determined at the trophic level.

*1.7. Effective biomass calculation*

We obtained body mass data for birds from EltonTraits (Wilman et al. 2014), while mass for arthropod species was estimated across the same representative species pools described above and using allometric equations per suborder, found predominantly in Sohlström et al. (2018), but also Hawkins et al. (1997; Gastropods), Hódar (1996; Acarina and Thysanoptera), and Townsend et al. (2012; Archaeognatha). For this purpose, body measurements for Coleoptera species (length and width) were taken from (Staab et al. 2025) and TraitCH (Chauvier-Mendes et al. 2026). For the remaining arthropods, we obtained body measurements (length) from various sources depending on taxon (Table S1). Where measurements were unavailable in any of these repositories, values were imputed manually using publicly available morphological descriptions and verified photographs from GBIF (GBIF.org 2026), processed with ImageJ (Schindelin et al. 2012) to extract standard body length measurements (Table S1).

For non-coleopteran arthropods, body mass was estimated for each representative species within each suborder (see above), and subsequently averaged per trophic level, weighted by diet proportion ($\bar{M_{ij}\cdot\alpha_{ijk}}$, where $\alpha_{ijk}$ is the species’ diet proportion at trophic level *k*, and $M_{ij}$ is the mass, per species *i*, in suborder *j*). This suborder- and trophic level-specific mass estimate was then multiplied by suborder abundance ($n_{j}$) to yield effective biomass per suborder and trophic level (${Be}_{jk}$). Effective biomass values were then summed across suborders to obtain total arthropod effective biomass at each trophic level (*k*):

$$arthropod {Be}_{k}=\sum_{j} (\bar{M_{ij}\cdot\alpha_{ijk}}){\cdot n}_{j}$$

To assess representativity of the suborders, the use of a species checklist was validated for Coleoptera 2023 data, where estimated checklist biomass was compared to the real biomass, with satisfactory results (Fig. S1). To evaluate whether the choice of species checklists used to derive suborder-level arthropod body mass estimates influenced our results, we conducted a sensitivity analysis in which all statistical models were repeated using the 25th and 75th percentile body mass estimates in place of the suborder mean. As results and interpretations were consistent across all three estimates, we report only the mean-based results. For coleopterans, biomass calculations were done at the species level (i.e. without averaging across subgroup) and added to the rest of the arthropods according to trophic level. Since Coleoptera were identified to species level in 2023 only, we applied the species proportions of each three-week sampling round from 2023 to the corresponding 2024 abundance data to estimate species composition, assuming that proportions in our study area remain stable between years under non-outbreak conditions (A. Szallies, personal communication).

Bird effective biomass was calculated using the same equation but at the species level, by multiplying the mass, diet proportion and abundance per species and summing the results for all species per plot and trophic level.

*1.8. Statistical analyses*

All statistical analyses were done in the R environment (version 4.5.1, R Core Team 2022). To identify the factors driving effective biomass and abundance, we scaled all continuous variables (mean = 0, SD = 1), enabling direct comparison of coefficient estimates as measures of relative importance (Schielzeth 2010). Scaling was applied to predictor and response variables, as response variables in one regression were used as predictors in subsequent trophic levels within the SEM. We assessed bivariate correlations among predictor variables (Table S2 and Fig. S2) and excluded variables with pairwise correlations exceeding r = 0.7. Where multiple correlated variables were candidates for inclusion, Principal Component Analyses (PCA) loading strengths informed selection of the most representative variable. Where correlated variables were theoretically relevant and could not be reduced to a single representative, PCAs were used to integrate them into a single dimension, which was only the case for microclimatic variables (temperature and relative humidity, mean and variability) treated separately for canopy and understorey layers. The first principal component (PC1) explained a large proportion of variance (75–87%) and was therefore retained as a composite microclimate metric (Figs. S3-S4). For mean microclimate variables, relative humidity was multiplied by −1 prior to analysis so that higher PC1 values correspond to warmer and drier conditions. The sign of PCA axes was adjusted where necessary to ensure consistent ecological interpretation without affecting statistical inference.

Global models included predictor variables from both layers regardless of the response variable’s layer, to allow assessment of across-layer effects (Table S4). To avoid overparameterization relative to sample size and improve model interpretability, we adopted a two-step approach in which predictor sets were further reduced using automated model selection using the ‘dredge’ function from the ‘MuMIn’ package (Bartoń 2010). Given the sample size (n = 33), predictor number was restricted to a maximum of seven to maintain a defensible ratio of observations to free parameters in a Bayesian framework and avoid overfitting (Lee and Song 2004). Model selection uncertainty was addressed by averaging coefficients across all candidate models with ΔAICc ≤ 2. Predictor variable support was quantified as the sum of Akaike weights across candidate models containing that variable (Burnham and Anderson 2002), and only variables with summed Akaike weights ≥ 0.5 were retained for subsequent structural equation modelling, as this threshold indicates a variable contributed to the majority of the cumulative model weight.

To investigate potential pathways by which structural vegetation attributes influence arthropod and bird abundance and biomass, we applied a piecewise Bayesian structural equation model (BSEM) for each measure separately (Fig. 1), using the ‘blavaan’ package (Merkle et al. 2015), with BSEM pathways following a bottom-up direction (Fig. 1). For the effective abundance model, residual covariances were manually specified between ecologically related variable pairs – canopy and understorey counts within arthropod trophic levels (n_arthropod_1_canopy ~~ n_arthropod_1_understorey, n_arthropod_2_canopy ~~ n_arthropod_2_understorey), between trophic levels within vertical layer (n_arthropod_1_canopy ~~ n_arthropod_2_canopy, n_arthropod_1_understorey ~~ n_arthropod_2_understorey), and between bird trophic levels (n_bird_1_total ~~ n_bird_2_total) – to account for shared unmeasured influences. Since posterior predictive p-values (PPP) values were satisfactory for biomass models, meaning that the implied covariance structure accurately reproduced the observed one, further path specification was not needed for these SEMs. We fitted models using default priors on regression paths (Normal (0, 10)), with initial MCMC values drawn within this range. To estimate models, we used four Markov chains, each with 8000 posterior samples following a 2000 sample burn-in phase. Model fit was evaluated using modification indices (MI) only after the first run, and additional paths were added when MI > 10 (Jones and Hofer 2016). We evaluated final model fit using PPP values, posterior predictive checks (comparing observed and simulated distributions), and R^2^ values. We assessed out-of-sample predictive performance and evaluated convergence of the MCMC chains using R̂ (R̂ < 1.1), effective sample size (ESS > 1000), Monte Carlo Standard Error (MCSE < 0.05) and visual inspection of chains. To further assess the robustness of our results, we repeated the statistical procedure using only Coleoptera, for which species-level identification provides the most reliable biomass estimates. Additionally, to test whether trophic links between arthropods and birds were obscured by taxa rarely consumed by birds (e.g. Acari, Thysanoptera), we repeated analyses restricting arthropods to groups known to dominate the diets of the occurring bird species (Coleoptera, Araneae, Diptera, Lepidoptera, Aphidina, Heteroptera and Holometabolic larvae). This resulted in six models, each assessing either abundance or biomass, for all arthropods, arthropods filtered by bird diet (hereafter referred to as filtered-arthropod model), or Coleoptera only.

2. RESULTS

Results are summarised in Fig. 2 and are otherwise here described in detail and refer to the specific panels per figure (Fig. 3, S6-S7).

Bottom-up biotic pathways showed vertical layer consistency across most models, where abundance and biomass of arthropod primary consumers at a specific layer consistently explained that of arthropod secondary consumers at the same layer (Fig. 3, S6-S7 panels C, D, I and J). Nevertheless, the direction of the effects on effective biomass and abundance differed among vegetation-related metrics; plant Rao’s Q in the understorey showed negative effects across arthropod predators (Fig. 3, S6-S7 panel C, Fig. S7 panel D), and a positive effect on herbivores in the understorey in the filtered arthropod and Coleoptera models (Figs. S6-S7 panel B). Rao’s Q in the understorey also showed positive effects on primary consumer birds (Fig. 3, S6-S7 panels E and K), while canopy plant Rao’s Q had consistent positive effects on arthropods (Fig. 3, S6 panel G, Fig. S6-S7 panel H). Plant species richness also showed generally negative effects on arthropod biomass and abundance (Fig. 3, S6-S7 panel D, Fig. 3 panel H, Fig. S6-S7 panel J), while plant cover in the canopy showed consistently negative effects across all arthropod trophic levels (Fig. 3, S6 panel D, Fig. 3, S7 panel H, Fig. 3 panel J, Fig. S7 panel B). PAI showed more complex patterns; while canopy PAI had negative effects on abundance and biomass across arthropod models (Fig. 3, S7 panel G, Fig. 3, panel H, Fig. 3, S6 panel I), understorey PAI showed mixed results, with positive effects on biomass across the arthropod biomass models (Fig. 3, S6 panel C, Fig. 3 panel F) and negative effects on the arthropod abundance models (Fig. S6-S7 panel H), although Coleoptera models presented exceptions to this pattern (Fig. S7 panels B and G). Further structural variables like canopy height showed largely negative effects on both effective biomass and abundance across all taxonomic groups (Fig. 3, S6-S7 panel D and L, Fig. 3, S7 panel B, Fig. 3 panel J, Fig. S7 panel H), while canopy height heterogeneity showed consistent positive effects across all models (Fig. 3, S6-S7 panel A and L, Fig. S7 panel D). Effective biomass and abundance were affected equally by microclimatic variables, with variability affecting all response variables positively across trophic levels (Fig. 3, S6-S7 panel K, Fig, S6-S7 panel I, Fig. S6 panel H, Fig. S7 panel D), while mean microclimate (high values meaning warm and dry) consistently showed negative effects on all arthropods (Fig. 3, S6 panels B and H, Fig. 3 panels D, I and J, Fig. S7 panel G) and positive effects on all birds (Fig. 3, S6-S7 panels F, K and L).

Brodbeck, Brent V., Joseph Funderburk, Julie Stavisky, Peter C. Andersen, and Jan Hulshof. 2002. ‘Recent Advances in the Nutritional Ecology of Thysanoptera, or the Lack Thereof’. *Thrips and Tospoviruses: Proceedings of the 7th International Symposium on Thysanoptera* (Canberra).

BugGuide (BugGuide). 2026. ‘BugGuide’. https://www.bugguide.net/.

Burnham, Kenneth P., and David R. Anderson, eds. 2002. *Model Selection and Multimodel Inference: A Practical Information-Theoretic Approach*. 2nd edn. Springer New York. https://doi.org/10.1007/b97636.

Chauvier-Mendes, Yohann, Antoine Adde, Ariel Bergamini, et al. 2026. ‘TraitCH: A Multi-Taxa Functional Trait Dataset for Switzerland and Europe’. Preprint, ESSD – Biosphere/Terrestrial biodiversity, February 3. https://doi.org/10.5194/essd-2025-754.

Cockroach Species File (Cockroach Species File). 2026. ‘Cockroach Species File’. http://cockroach.archive.speciesfile.org/HomePage/Cockroach/HomePage.aspx.

Dengler, Jürgen, and Iwona Dembicz. 2023. ‘Should We Estimate Plant Cover in Percent or on Ordinal Scales?’ *Vegetation Classification and Survey* 4: 131–38. https://doi.org/10.3897/VCS.98379.

DGfO (DGfO). 2026. ‘Deutsche Gesellschaft Für Orthopterologie’. https://dgfo-articulata.de/schaben.

Díaz, Sandra, Jens Kattge, Johannes H. C. Cornelissen, et al. 2016. ‘The Global Spectrum of Plant Form and Function’. *Nature* 529 (7585): 167–71. https://doi.org/10.1038/nature16489.

Doğanlar, Mikdat. 2018. ‘Species of Macroglenes WESTWOOD, 1832 (Hymenoptera: Pteromalidae, Spalangiinae) from Turkey, with Description of a New Species’. *Entomofauna* 39 (14): 317–24.

Duelli, Peter, and Charles S. Henry. 2022. ‘The Apertochrysa Prasina Group (Neuroptera: Chrysopidae), with a Key to the European Species’. *Zootaxa* 5134 (1): 61–91. https://doi.org/10.11646/zootaxa.5134.1.3.

Falkner, Gerhard, Petr Obrdlík, Emmanuel Castella, and Martin CD Speight. 2001. *Shelled Gastropoda of Western Europe*. Friedrich-Helds Gesellschaft.

FOEN, Federal Office for the Environment and InfoSpecies. 2023. *Gefährdete Arten und Lebensräume in der Schweiz: Synthese Rote Listen*. Bern.

Garcia-Romera, Carlos, and Jose A. Barrientos. 2017. ‘Structure of Scuttle Fly Communities (Diptera: Phoridae) in Two Habitats on a Mediterranean Mountain’. *European Journal of Entomology* 114 (April): 203–14. https://doi.org/10.14411/eje.2017.025.

GBIF.org (GBIF Home Page). 2026. ‘GBIF Backbone Taxonomy. Checklist Dataset’. https://doi.org/10.15468/39omei.

Gossner, Martin M., Nadja K. Simons, Roland Achtziger, et al. 2015. ‘A Summary of Eight Traits of Coleoptera, Hemiptera, Orthoptera and Araneae, Occurring in Grasslands in Germany’. *Scientific Data* 2 (1). https://doi.org/10.1038/sdata.2015.13.

Gracillariidae.Net (Gracillariidae.Net). 2026. ‘Gracillariidae.Net’. https://www.gracillariidae.net/.

Haas, Michael, Hannes Baur, Tanja Schweizer, et al. 2021. ‘Tiny Wasps, Huge Diversity – A Review of German Pteromalidae with New Generic and Species Records (Hymenoptera: Chalcidoidea)’. *Biodiversity Data Journal* 9 (December): e77092. https://doi.org/10.3897/BDJ.9.e77092.

Harrison, Douglas A., and Robin L. Cooper. 2003. ‘Characterization of Development, Behavior and Neuromuscular Physiology in the Phorid Fly, Megaselia Scalaris’. *Comparative Biochemistry and Physiology Part A: Molecular & Integrative Physiology* 136 (2): 427–39. https://doi.org/10.1016/S1095-6433(03)00200-9.

Hawkins, J. W., M. W. Lankester, R. A. Lautenschlager, and F. W. Bell. 1997. ‘Length–Biomass and Energy Relationships of Terrestrial Gastropods in Northern Forest Ecosystems’. *Canadian Journal of Zoology* 75 (3): 501–5. https://doi.org/10.1139/z97-061.

Heckmann, Ralf, and Hermann Blöchlinger. 2011. *Die Wanzenfauna (Hemiptera: Heteroptera) des Kantons Thurgau, Teil 1. Dipsocoromorpha, Nepomorpha, Gerromorpha, Leptopodomorpha und Cimicomorpha*. Text/html,application/pdf,text/html. https://doi.org/10.5169/SEALS-593838.

Hiermann, Ulrich. 2025. ‘Notizen zur Entomofauna Liechtensteins (Insecta: Archaeognatha, Plecoptera, Blattodea, Orthoptera, Heteroptera, Hymenoptera)’. *Inatura* 132 (7).

Hódar, José A. 1996. ‘The Use of Regression Equations for Estimation of Arthropod Biomass in Ecological Studies’. *Acta Oecologica* 17 (5): 421–33.

Hoebeke, E. Richard, and Maureen E. Carter. 2010. ‘First North American Record of *Ectobius Lucidus* (Hagenbach) (Blattodea: Blattellidae: Ectobiinae), with Notes on Recognition Characters and Seasonal History, and Additional Records for Other *Ectobius* Species in the Northeastern United States’. *Proceedings of the Entomological Society of Washington* 112 (2): 229–38. https://doi.org/10.4289/0013-8797-112.2.229.

InfluentialPoints (InfluentialPoints). 2026. ‘InfluentialPointsDotCom’. https://influentialpoints.com/Index.htm.

Info Fauna. 2026. ‘info fauna – National Data and Information Center for the Swiss Fauna’. Switzerland. https://www.infofauna.ch/de#gsc.tab=0.

Infusino, Marco, and Stefano Scalercio. 2018. ‘The Importance of Beech Forests as Reservoirs of Moth Diversity in Mediterranean Basin (Lepidoptera)’. *Fragmenta Entomologica* 50 (2): 161–70. https://doi.org/10.4081/fe.2018.294.

Insects.CH (Insects.CH). 2026. ‘Insects.CH’. https://www.insects.ch/.

Insekten Sachs. (Insekten Sachsen). 2026. ‘Insekten Sachsen’. https://www.insekten-sachsen.de.

Isenburg, M. 2024. *LASTools - Efficient LiDAR Processing Software*. V. 241125. Rapidlasso GmbH, released. https://rapidlasso.com/lastools/.

Jones, N., and Scott M. Hofer. 2016. *Structural Equation Modeling with AMOS: Basic Concepts, Applications, and Programming*. 3rd edn. New York. https://doi.org/https://doi.org/10.4324/9781315757421.

Käfer Eur. (Die Käfer Europas). 2026. ‘Die Käfer Europas’. https://coleonet.de/coleo/index.htm.

Kattge, Jens, Gerhard Bönisch, Sandra Díaz, et al. 2020. ‘TRY Plant Trait Database – Enhanced Coverage and Open Access’. *Global Change Biology* 26 (1): 119–88. https://doi.org/10.1111/gcb.14904.

Keilin, D. 1940. ‘THE EARLY STAGES OF THE FAMILES TRICHOCERIDAE AND ANISOPODIDE (=RHYPHIDAE)(DIPTERA: NEMATOCERA)’. *Transactions of the Royal Entomological Society of London* 90 (3): 39–62. https://doi.org/10.1111/j.1365-2311.1940.tb00742.x.

Klopfstein, Seraina, Matthias Riedel, and Martin Schwarz. 2019. ‘Checklist of Ichneumonid Parasitoid Wasps in Switzerland (Hymenoptera, Ichneumonidae): 470 Species New for the Country and an Appraisal of the Alpine Diversity’. *Alpine Entomology* 3 (April): 51–81. https://doi.org/10.3897/alpento.3.31613.

Kowalski, Esther, Martin M. Gossner, Manfred Türke, et al. 2011. ‘The Use of Forest Inventory Data for Placing Flight-Interception Traps in the Forest Canopy’. *Entomologia Experimentalis et Applicata* 140 (1): 35–44. https://doi.org/10.1111/j.1570-7458.2011.01134.x.

Kucharczyk, Halina, and Marek Kucharczyk. 2013. ‘Characteristic and Diagnostic Features of the Most Frequently Occurring Species of the Thripidae Family (Insecta, Thysanoptera) in Crown Canopies of Central European Forests’. *Forest Research Papers* 74 (1): 5–11. https://doi.org/10.2478/frp-2013-0001.

Kükenbrink, Daniel, Mauro Marty, Nataliia Rehush, Meinrad Abegg, and Christian Ginzler. 2025. ‘Evaluating the Potential of Handheld Mobile Laser Scanning for an Operational Inclusion in a National Forest Inventory – A Swiss Case Study’. *Remote Sensing of Environment* 321 (May): 114685. https://doi.org/10.1016/j.rse.2025.114685.

Külling, Nathan, Antoine Adde, Fabian Fopp, et al. 2024. ‘SWECO25: A Cross-Thematic Raster Database for Ecological Research in Switzerland’. *Scientific Data* 11 (1). https://doi.org/10.1038/s41597-023-02899-1.

Laliberté, Etienne, Pierre Legendre, and Bill Shipley. 2023. *FD: Measuring Functional Diversity (FD) from Multiple Traits, and Other Tools for Functional Ecology*. V. 1.0-12.3. Released November 26. https://cran.r-project.org/web/packages/FD/index.html.

Landolt, E., B. Bäumler, A. Erhardt, et al. 2010. *Flora Indicativa. Ökologische Zeigerwerte Und Biologische Kennzeichen Zur Flora Der Schweiz Und Der Alpen. Ecological Indicators Values and Biological Attributes of the Flora of Switzerland and the Alps.* Haupt.

Lee, Sik-Yum, and Xin-Yuan Song. 2004. ‘Evaluation of the Bayesian and Maximum Likelihood Approaches in Analyzing Structural Equation Models with Small Sample Sizes’. *Multivariate Behavioral Research* 39 (4): 653–86. https://doi.org/10.1207/s15327906mbr3904_4.

LepiWiki (LepiWiki). 2026. ‘LepiWiki’. https://lepiforum.org/wiki.

Lithocolletinae North-West Eur. (Lithocolletinae of North-West Europe). 2026. ‘Lithocolletinae of North-West Europe’. https://lithocolletinae.linnaeus.naturalis.nl/linnaeus_ng/app/views/matrixkey/index.php?epi=9.

Martel, Jocelyn, and Yves Mauffette. 1997. ‘Lepidopteran Communities in Temperate Deciduous Forests Affected by Forest Decline’. *Oikos* 78 (1): 48. https://doi.org/10.2307/3545799.

Merkle, Edgar, Yves Rosseel, and Ben Goodrich. 2015. *Blavaan: Bayesian Latent Variable Analysis*. Released October 7. https://CRAN.R-project.org/package=blavaan.

Morsdorf, Felix, Mauro Marty, and Daniel Kükenbrink. 2025. ‘UAV-Based LiDAR and SfM-Derived 3D Point Clouds of Forest Canopies - Observation Angles Matter’. *Dreiländertagung D-A-CH 2025 ‘Raumbezogene Bilddaten Und Künstliche Intelligenz Für Nachhaltige Lebensräume’*, 338–50. Application/pdf. https://doi.org/10.24407/KXP:1928725848.

Mühlethaler, Roland, Valeria Trivellone, Roel van Klink, Rolf Niedringhaus, and Herbert Nickel. 2016. ‘Kritische Artenliste der Zikaden der Schweiz (Hemiptera: Auchenorrhyncha)’. *Cicadina* 16 (4): 49–87.

Murphy, Stephen J., André M. Bellvé, Reymond J. Miyajima, et al. 2023. ‘SAviTraits 1.0: Seasonally Varying Dietary Attributes for Birds’. *Global Ecology and Biogeography* 32 (10): 1690–98. https://doi.org/10.1111/geb.13738.

NatureSpot (NatureSpot). 2026. ‘NatureSpot’. https://www.naturespot.org/.

Pimont, François, Denis Allard, Maxime Soma, and Jean-Luc Dupuy. 2018. ‘Estimators and Confidence Intervals for Plant Area Density at Voxel Scale with T-LiDAR’. *Remote Sensing of Environment* 215 (September): 343–70. https://doi.org/10.1016/j.rse.2018.06.024.

Planthoppers Psyllist (Planthoppers: Psyllist). 2026. ‘Planthoppers: Psyllist’. http://flow.hemiptera-databases.org/psyllist/.

R Core Team. 2025. *R: A Language and Environment for Statistical Computing*. Vienna, Austria, Released. https://www.R-project.org/.

Schielzeth, Holger. 2010. ‘Simple Means to Improve the Interpretability of Regression Coefficients’. *Methods in Ecology and Evolution* 1 (2): 103–13. https://doi.org/10.1111/j.2041-210x.2010.00012.x.

Schindelin, Johannes, Ignacio Arganda-Carreras, Erwin Frise, et al. 2012. ‘Fiji: An Open-Source Platform for Biological-Image Analysis’. *Nature Methods* 9 (7): 676–82. https://doi.org/10.1038/nmeth.2019.

Schneider, Fabian D., Daniel Kükenbrink, Michael E. Schaepman, David S. Schimel, and Felix Morsdorf. 2019. ‘Quantifying 3D Structure and Occlusion in Dense Tropical and Temperate Forests Using Close-Range LiDAR’. *Agricultural and Forest Meteorology* 268 (April): 249–57. https://doi.org/10.1016/j.agrformet.2019.01.033.

Schneider, Fabian D., Felix Morsdorf, Bernhard Schmid, et al. 2017. ‘Mapping Functional Diversity from Remotely Sensed Morphological and Physiological Forest Traits’. *Nature Communications* 8 (1). https://doi.org/10.1038/s41467-017-01530-3.

Sohlström, Esra H., Lucas Marian, Andrew D. Barnes, et al. 2018. ‘Applying Generalized Allometric Regressions to Predict Live Body Mass of Tropical and Temperate Arthropods’. *Ecology and Evolution* 8 (24): 12737–49. https://doi.org/10.1002/ece3.4702.

Staab, Michael, Nadja Simons, and Martin Goßner. 2025. ‘Body Size and Life-History Traits of Arthropod Species’. Version 18. Biodiversity Exploratories Information System. www.bexis.uni-jena.de/ddm/data/Showdata/31122?tag=18.

Sturm, Helmut. 1996. ‘Die Machiliden (Archaeognatha, Apterygota, Insecta) Nord westdeutschlands und die tiergeographische Bedeutung dieser Vorkommen’. *Drosera* 80 (2): 53–62.

Sven. Fjärilar NRM (Svenska Fjärilar NRM). 2014. ‘Svenska Fjärilar - Naturhistoriska Riksmuseet’. http://www3.nrm.se/en/svenska_fjarilar/svenska_fjarilar.html.

swisstopo. 2023. ‘Waldreservate Der Schweiz’. Federal Office for the Environment FOEN. https://www.geocat.ch/geonetwork/srv/eng/catalog.search#/metadata/4b9ff750-3e52-43e9-80d1-c07c59c469fd.

Szikora, Timea. 2015. ‘Promoting Parasitic Wasps among Swiss Lowland Extensively Managed Meadows: Positive Effects of Delaying Mowing and Leaving Uncut Grass Refuges’. MSc, Universität Bern.

Tauber, Catherine A., Maurice J. Tauber, and Gilberto S. Albuquerque. 2009. ‘Neuroptera’. In *Encyclopedia of Insects*. Elsevier. https://doi.org/10.1016/B978-0-12-374144-8.00190-9.

Todorov, Ivaylo, Mircea-Dan Mitroiu, Aneliya Bobeva, and Peter Boyadzhiev. 2024. ‘Description of Mesopolobus Askewi Sp. Nov. (Hymenoptera, Pteromalidae), with Notes on the Fauna of Asaphesinae and Pteromalidae (Hymenoptera, Chalcidoidea) Collected from Foliage of Picea Abies (L.) H. Karst. in Bulgaria’. *Biodiversity Data Journal* 12 (December): e139403. https://doi.org/10.3897/BDJ.12.e139403.

Townsend, Jason M., Christopher C. Rimmer, Kent P. McFarland, and James E. Goetz. 2012. ‘Site-Specific Variation in Food Resources, Sex Ratios, and Body Condition of an Overwintering Migrant Songbird’. *The Auk* 129 (4): 683–90. https://doi.org/10.1525/auk.2012.12043.

Ulrich, W., and K. Szpila. 2008. ‘Body Size Distributions of Eastern European Diptera’. *Polish Journal of Ecology* 56 (4): 557–68.

Vincent, Grégoire, Cécile Antin, Marilyne Laurans, et al. 2017. ‘Mapping Plant Area Index of Tropical Evergreen Forest by Airborne Laser Scanning. A Cross-Validation Study Using LAI2200 Optical Sensor’. *Remote Sensing of Environment* 198 (September): 254–66. https://doi.org/10.1016/j.rse.2017.05.034.

Wilman, Hamish, Jonathan Belmaker, Jennifer Simpson, Carolina De La Rosa, Marcelo M. Rivadeneira, and Walter Jetz. 2014. ‘EltonTraits 1.0: Species‐level Foraging Attributes of the World’s Birds and Mammals: Ecological Archives E095‐178’. *Ecology* 95 (7): 2027–2027. https://doi.org/10.1890/13-1917.1.

3. FIGURES

Figure 1. Schematic depiction of the global models used to inform the regressions included in the Bayesian SEM. Paths from each explanatory variable (here grouped into microclimatic, plant diversity and vegetation structure, right) were directly linked to each response variable within the green circle (repeated for abundance and biomass separately for the four different trophic and taxonomic groups, left). Explanatory variables of different vertical layers were allowed to coexist in the global models, prior to automated model selection and imputation into the SEM. Nevertheless, canopy height and canopy height heterogeneity were defined at the plot-level, not at the layer-level. Global models for taxonomic groups of trophic level two included the relevant response variables at trophic level one as explanatory variables. For example, this was the case for the global model of bird abundance or biomass for trophic level two (warbler silhouette, top centre), which included all explanatory variables outside the green circle as well as invertebrate (abundance or biomass respectively) at trophic level one (thrips silhouette bottom right) and two (ladybug silhouette centre left).

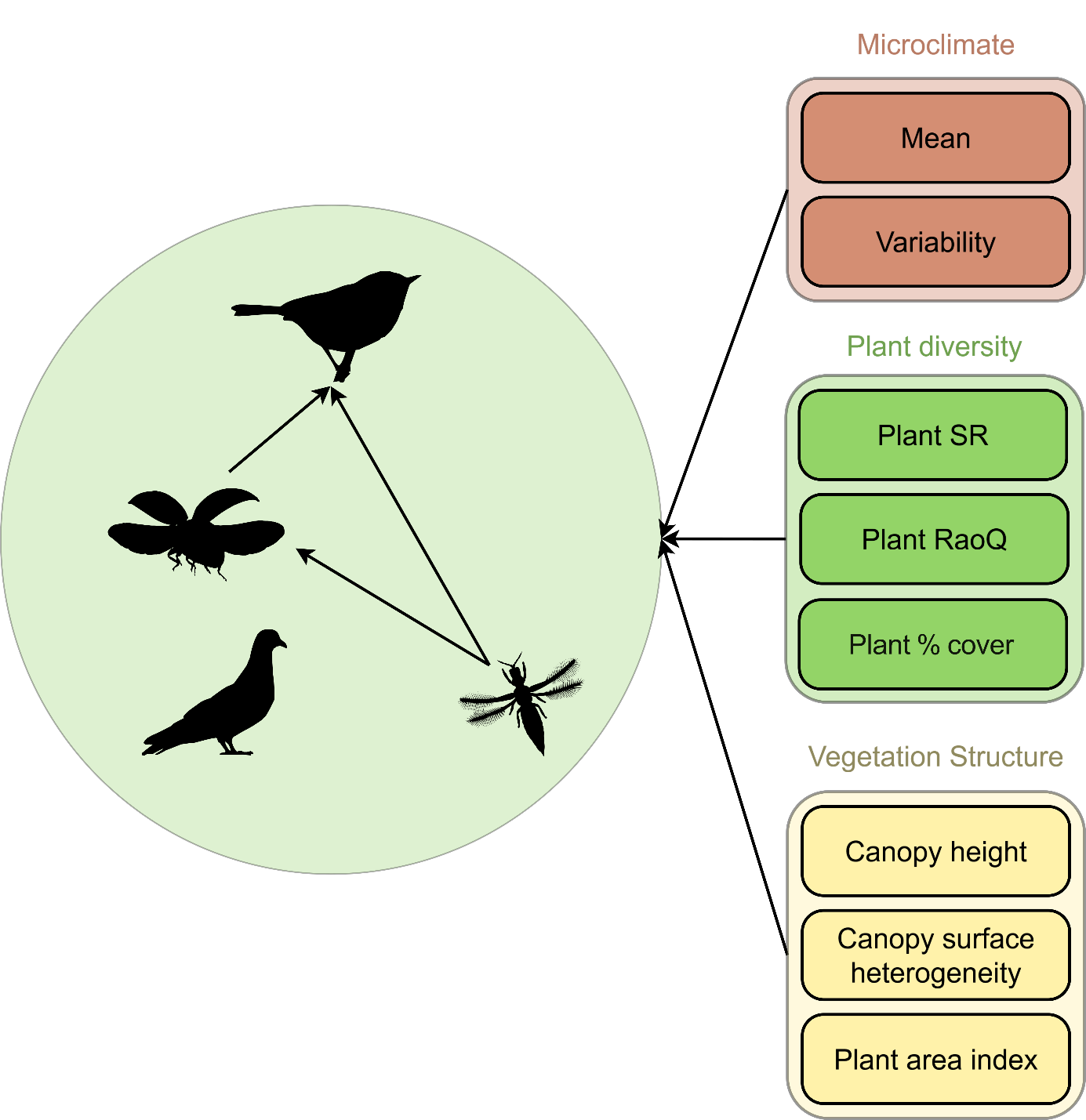

| Taxon | Repr. sp. | Checklist scale | Checklist source | Diet data source | Length measurement source |
| --- | --- | --- | --- | --- | --- |
| Acarina | 5 | Study region + Europe | Info Fauna 2026; NatureSpot 2026 | NatureSpot 2026 | NatureSpot 2026 |
| Aphidina | 21 | Europe | InfluentialPoints 2026 | Auclair 1963 | InfluentialPoints 2026 |
| Apocrita | 18 | Study region + Switzerland | Info Fauna 2026; Szikora 2015; Klopfstein et al. 2019 | Gossner et al. 2015; Chauvier-Mendes et al. 2026 | Gossner et al. 2015; Chauvier-Mendes et al. 2026; Haas et al. 2021; Baur et al. 2007; Todorov et al. 2024; Doğanlar 2018 |
| Araneae | 56 | Study region | Info Fauna 2026 | Gossner et al. 2015; Chauvier-Mendes et al. 2026 | Gossner et al. 2015; Chauvier-Mendes et al. 2026 |
| Archaeognatha | 3 | Central Europe | Hiermann 2025; Sturm 1996 | Sturm 1996 | Sturm 1996; GBIF.org 2026 |
| Blattodea | 3 | Central Europe | DGfO 2026; Cockroach Species File 2026 | Insekten Sachs. 2026; BugGuide 2026; Hoebeke and Carter 2010 | Insekten Sachs. 2026; BugGuide 2026; Hoebeke and Carter 2010 |
| Brachycera | 45 | Study region | Info Fauna 2026 | expert knowledge at family level | Ulrich and Szpila 2008 |
| Chilopoda | 1 | Central Europe | ‘Bodentier^4^’ 2026 | Bodentier^4^ 2026 | Bodentier^4^ 2026 |
| Cicadina | 11 | Switzerland | Mühlethaler et al. 2016 | Gossner et al. 2015 | Gossner et al. 2015 |
| *Coleoptera* | *570* | *-* | *-* | *Gossner et al. 2015; Chauvier-Mendes et al. 2026* | *Gossner et al. 2015; Chauvier-Mendes et al. 2026; Käfer Europas 2026; GBIF.org 2026* |
| Dermaptera | 3 | Central Europe | DGfO 2026 | Insekten Sachs. 2026 | Insekten Sachs. 2026 |
| Ensifera | 5 | Study region | Info Fauna 2026 | Gossner et al. 2015; Chauvier-Mendes et al. 2026 | Gossner et al. 2015; Chauvier-Mendes et al. 2026 |
| Formicidae | 5 | Study region | Info Fauna 2026 | AntWiki 2026 | AntWiki 2026 |
| Gastropoda | 15 | Switzerland | Info Fauna 2026 | Info Fauna 2026 | Falkner et al. 2001 |
| Heteroptera | 133 | Study region | Heckmann and Blöchlinger 2011 | Gossner et al. 2015 | Gossner et al. 2015 |
| Larvae-holo | 5 | Central Europe | Martel and Mauffette 1997; Infusino and Scalercio 2018; Böttger et al. 2025; Garcia-Romera and Barrientos 2017 | Sven. Fjärilar NRM 2014; Keilin 1940; Harrison and Cooper 2003 | Sven. Fjärilar NRM 2014; Keilin 1940; Harrison and Cooper 2003 |
| Lepidoptera | 40 | Study region | Info Fauna 2026; Sven. Fjärilar NRM 2014; LepiWiki 2026; Lithocolletinae North-West Eur. 2026; Gracillariidae.Net 2026 | Sven. Fjärilar NRM 2014; LepiWiki 2026; Lithocolletinae North-West Eur. 2026; Gracillariidae.Net 2026 | GBIF.org 2026; LepiWiki 2026; Gracillariidae.Net 2026; Lithocolletinae North-West Eur. 2026 |
| Nematocera | 19 | Study region | Info Fauna 2026 | expert knowledge at family level | Ulrich and Szpila 2008 |
| Neuroptera | 16 | Study region | Info Fauna 2026; Duelli and Henry 2022 | Tauber et al. 2009 | GBIF.org 2026 |
| Opiliones | 16 | Switzerland + Central Europe | Gossner et al. 2015, Chauvier-Mendes et al. 2026 | Gossner et al. 2015, Chauvier-Mendes et al. 2026 | Gossner et al. 2015, Chauvier-Mendes et al. 2026 |
| Pseudoscorpiones | 5 | Switzerland | Chauvier-Mendes et al. 2026 | Chauvier-Mendes et al. 2026 | Chauvier-Mendes et al. 2026 |
| Psocoptera | 3 | Central Europe | Insects.CH 2026 | Berry et al. 2008 | Insects.CH 2026 |
| Psyllidae | 7 | Central Europe | Planthoppers Psyllist 2026 | Planthoppers Psyllist 2026 | GBIF.org 2026 |
| Symphyta | 16 | Central Europe | Gossner et al. 2015 | Gossner et al. 2015 | Gossner et al. 2015 |
| Thysanoptera | 6 | Central Europe | Kucharczyk and Kucharczyk 2013 | Brodbeck et al. 2002 | GBIF.org 2026 |

Table S1. General information on the species checklists used per suborder, showing number of representative species (Repr. sp.) per group, spatial scale at which the checklist data was available, and source for the checklist, as well as the source for the diet and length information of the checklist species. No checklist was used for Coleoptera (italics).

| Predictor variable | | Description | Resolution | Mean | SD |
| --- | --- | --- | --- | --- | --- |
| *Microclimatic* | |  |  |  |  |
|  | microclim_m | PC1 of PCA for mean temperature (°C) and mean relative humidity (%), taken as the mean hourly values across the sampling period (April-June). High values indicate warm and dry environments | layer | - | - |
|  | microclim_var | PC1 of PCA for temperature and relative humidity standard deviation, taken as the SD of the hourly values across the sampling period (April-June). High values indicate high microclimate variability | layer | - | - |
| *Vegetation structure* | |  |  |  |  |
|  | CH_m | Mean canopy height (m) of the plot area calculated from the canopy height model defined as the 99th height percentile at 1 m resolution | plot | 22.81 | 6.49 |
|  | CH_sd | Canopy height standard deviation (heterogeneity) of the plot area calculated from the canopy height model defined as the 99th height percentile at 1 m resolution | plot | 4.80 | 2.96 |
|  | PAI_leafoff | Plant Area index calculated from leaf off acquisition data; measure of all plant parts per plot area | canopy | 4.32 | 1.91 |
|  |  |  | understorey | 0.37 | 0.19 |
| *Biotic* |  |  |  |  |  |
|  | plant_raoQ | Woody plant Rao's Q using percentage cover from the vegetation survey and traits gathered from the TRY database | canopy | 0.01 | 0.01 |
|  |  |  | understorey | 0.01 | 0.01 |
|  | plant_sr | Woody plant species richness according to the vegetation survey | canopy | 3.61 | 1.61 |
|  |  |  | understorey | 4.00 | 2.37 |
|  | cover_plant | Woody plant percentage cover according to the vegetation survey | canopy | 75.52 | 13.32 |
|  |  |  | understorey | 14.94 | 16.83 |
|  | forest_type | Plot categorical stand type (i.e. coniferous or broadleaf, as > 50 %), taken from canopy percentage cover according to the vegetation survey | plot | - | - |

Table S2. Predictor variable labels and descriptions used in our models. The resolution at which they were available (at the layer- (canopy and understorey) and plot-level), and their mean and SD are also shown.

| Taxon | Total ind. | Representative sp. | Mean Length (mm) | SD Length (mm) | Mean Mass (mg) | SD Mass (mg) | Mean Diet Proportion 1 | Mean Diet Proportion 2 |
| --- | --- | --- | --- | --- | --- | --- | --- | --- |
| Acarina | 9729 | 5 | 1.78 | 0.00 | 0.51 | 0.74 | 0.20 | 0.80 |
| Aphidina | 7771 | 21 | 2.62 | 0.00 | 1.63 | 1.28 | 1.00 | 0.00 |
| Apocrita | 2673 | 18 | 5.74 | 0.05 | 24.61 | 46.74 | 0.28 | 0.72 |
| Araneae | 431 | 56 | 4.39 | 0.01 | 10.77 | 7.30 | 0.00 | 1.00 |
| Archaeognatha | 2 | 3 | 10.08 | 0.00 | 7.08 | 3.96 | 1.00 | 0.00 |
| Blattodea | 37 | 3 | 9.22 | 0.01 | 30.31 | 5.39 | 0.00 | 1.00 |
| Brachycera | 5452 | 45 | 8.36 | 0.01 | 21.61 | 13.71 | 0.90 | 0.10 |
| Chilopoda | 1 | 1 | 20.00 | - | 88.28 | - | 0.00 | 1.00 |
| Cicadina | 362 | 11 | 5.48 | 0.00 | 10.13 | 4.17 | 1.00 | 0.00 |
| *Coleoptera* | *50080* | *570* | *4.48* | *0.03* | *11.02* | *34.32* | *0.23* | *0.55* |
| Dermaptera | 12 | 3 | 13.33 | 0.01 | 48.53 | 10.49 | 0.00 | 1.00 |
| Ensifera | 5 | 5 | 13.83 | 0.04 | 92.27 | 35.38 | 0.43 | 0.57 |
| Formicidae | 408 | 5 | 4.38 | 0.00 | 2.97 | 1.23 | 0.00 | 1.00 |
| Gastropoda | 19 | 15 | 8.01 | 0.01 | 32.78 | 11.56 | 0.40 | 0.27 |
| Heteroptera | 5590 | 133 | 6.11 | 0.02 | 12.67 | 15.01 | 0.60 | 0.39 |
| Larven-Holo | 4765 | 5 | 11.86 | 0.03 | 27.20 | 30.21 | 0.60 | 0.00 |
| Lepidoptera | 939 | 40 | 13.44 | 0.03 | 42.35 | 33.64 | 0.93 | 0.08 |
| Nematocera | 9388 | 19 | 7.22 | 0.02 | 10.59 | 18.98 | 0.47 | 0.37 |
| Neuroptera | 732 | 16 | 5.71 | 0.00 | 6.60 | 2.76 | 0.00 | 1.00 |
| Opiliones | 11 | 16 | 4.47 | 0.01 | 16.07 | 7.29 | 0.23 | 0.35 |
| Pseudoscorpiones | 7 | 5 | 2.66 | 0.00 | 0.91 | 0.62 | 0.00 | 1.00 |
| Psocoptera | 1450 | 3 | 2.02 | 0.00 | 0.55 | 0.47 | 1.00 | 0.00 |
| Psyllidae | 670 | 7 | 2.57 | 0.00 | 1.33 | 0.44 | 1.00 | 0.00 |
| Symphyta | 155 | 16 | 8.90 | 0.02 | 33.99 | 22.66 | 1.00 | 0.00 |
| Thysanoptera | 29400 | 6 | 2.02 | 0.00 | 0.04 | 0.02 | 1.00 | 0.00 |

Table S3. Summary information for the suborder groups using the checklist method (except for Coleoptera, in italics, species level data shown). The total captured individuals and number of representative species used for the checklist pool are shown, as well as the mean and SD for length and mass for these checklists’ species. The proportion of species in trophic level one and two according to the checklists (mean diet proportion 1 and 2) does not necessarily add up to one since decomposers were not included in the study.

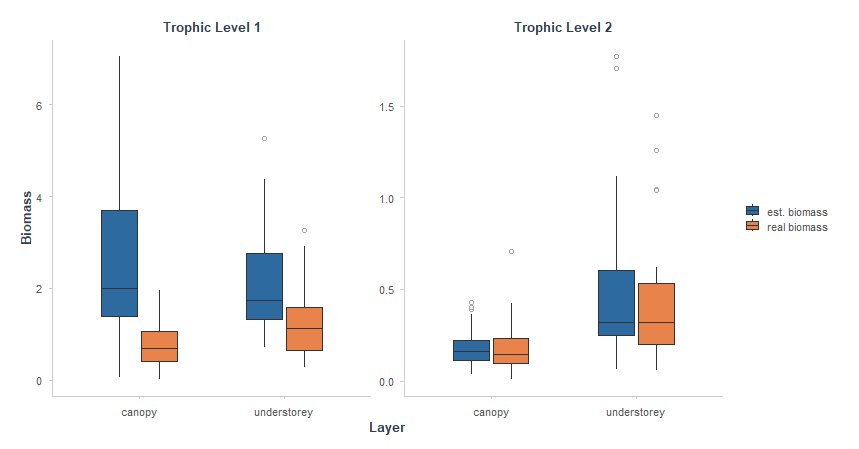

Figure S1. Comparison between estimated biomass (using the checklist method, blue) and real biomass (orange) for coleoptera data of 2023. Trophic level one (23% of coleoptera, see Table S3) shows an overestimation when using estimated biomass for this taxonomic group, while trophic level two shows similar results between both biomass values. This difference between estimates and real values in trophic level one supported the further sensitivity analysis during the BSEM procedure.

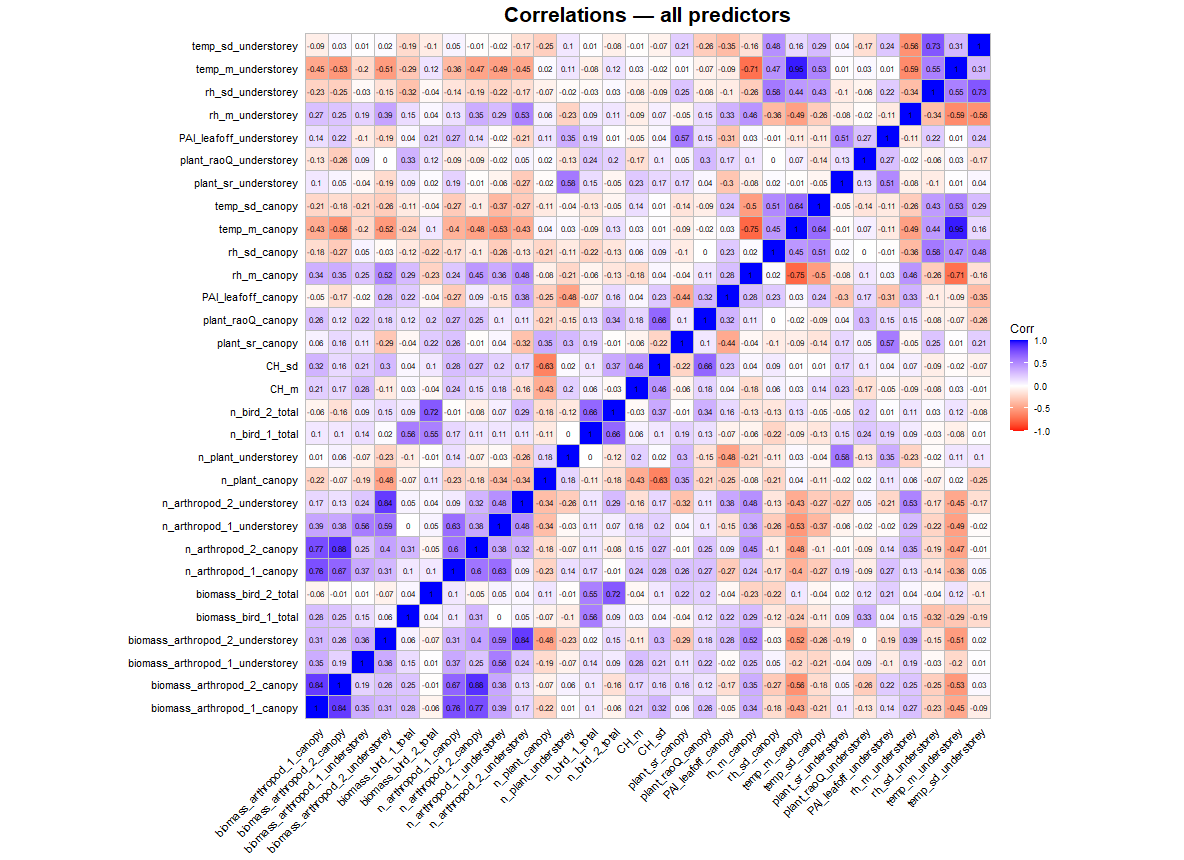

Figure S2. Pearson’s bivariate correlation plot for all continuous response and predictor variables used in the SEMs. Effective abundance variables are labelled as ‘n_’, while the trophic level is marked with 1 or 2 for primary and secondary consumers. Microclimatic variables were thereafter reduced using PCAs due to their high correlation coefficients.

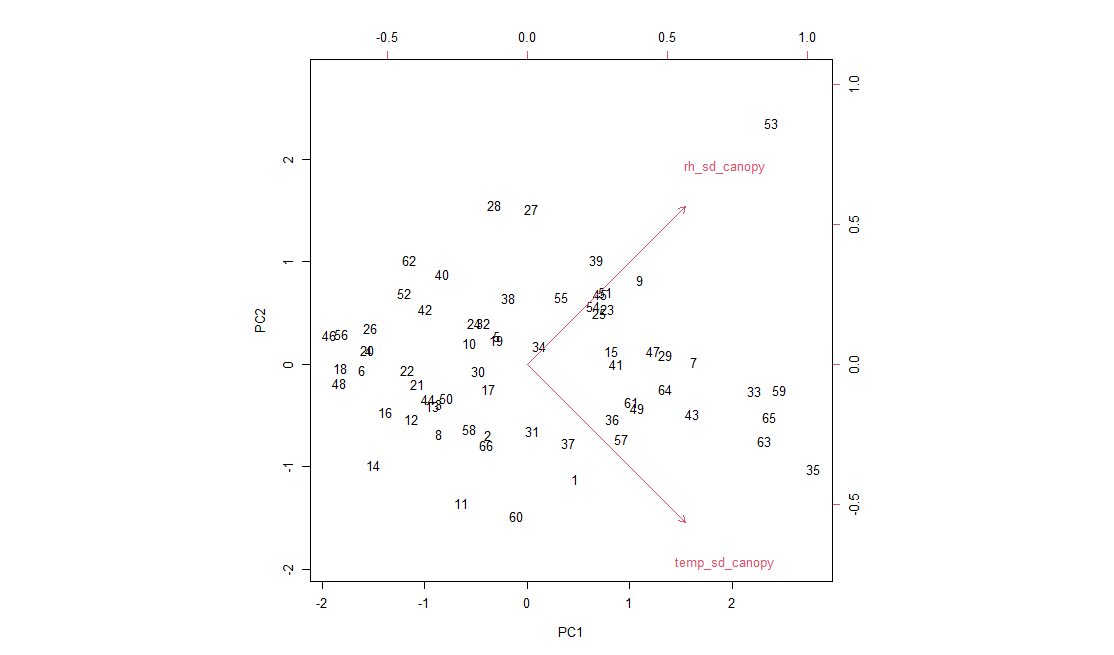

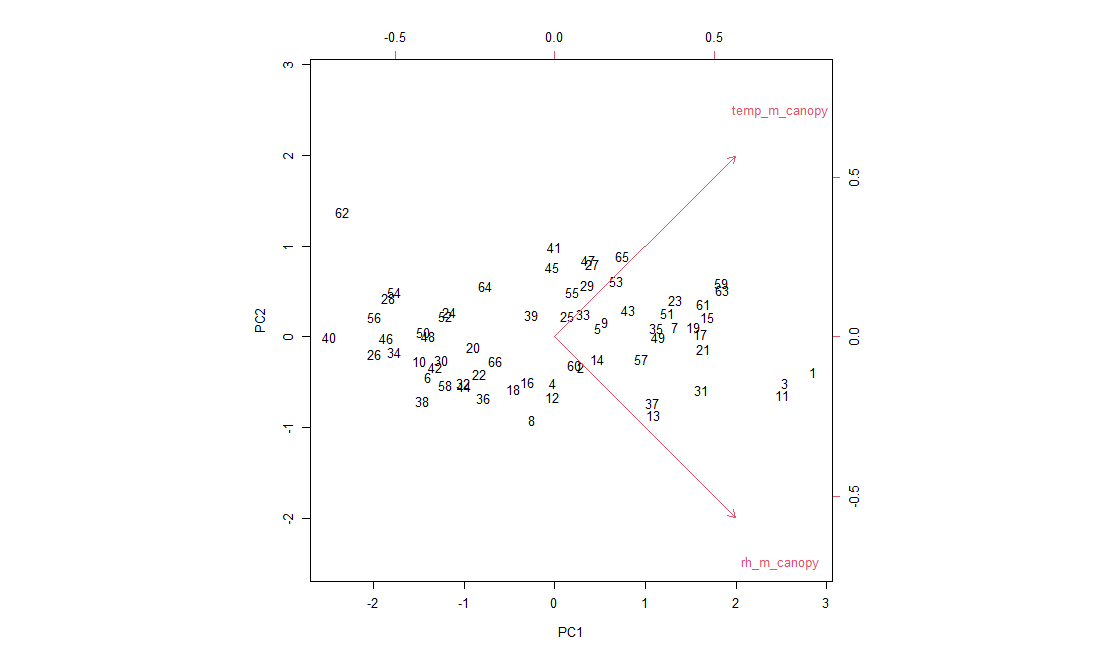

Figure S3. Partial Components Analysis (PCA) plot combining microclimatic predictor variables for the mean (above) and standard deviation (below) temperature and relative humidity measurements at the canopy layer. The PC1 only was selected (0.87 for mean microclimate and 0.75 for microclimate variability). The loadings are inverted to become positive (higher microclimatic ‘mean’ or ‘variability’ means higher PC1 values).

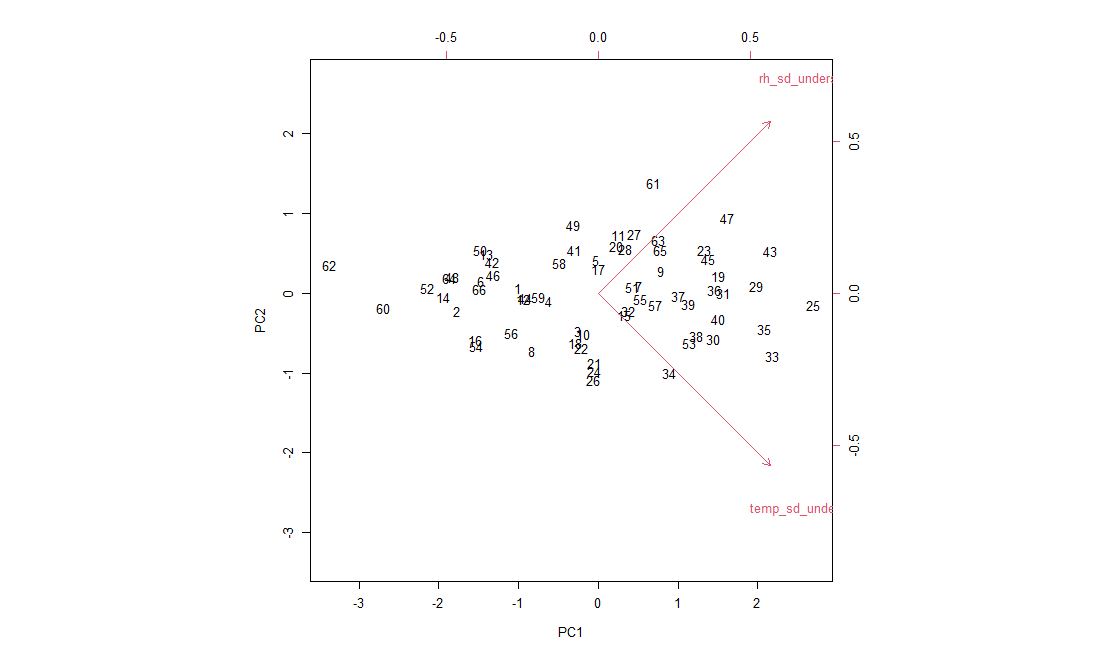

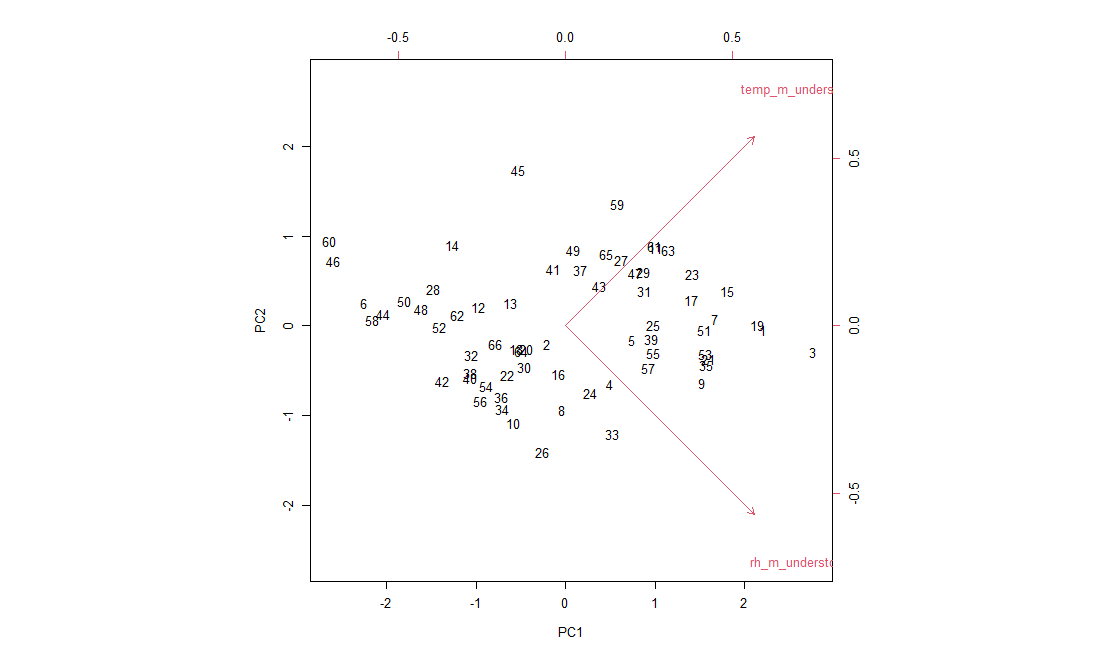

Figure S4. Partial Components Analysis (PCA) plot combining microclimatic predictor variables for the mean (above) and standard deviation (below) temperature and relative humidity measurements at the understorey layer. The PC1 only was selected (0.79 for mean microclimate and 0.86 for microclimate variability). The loadings are inverted to become positive (higher microclimatic ‘mean’ or ‘variability’ means higher PC1 values).

Table S4. Structure of global models prior to automated variable selection. For each response variable, the full set of fixed effects included in the model is shown. Global model structures were identical for effective biomass and abundance responses (only effective biomass models are presented here).

| Response | Predictors |
| --- | --- |
| biomass_arthropod_1_canopy | year + plant_sr_canopy + plant_sr_understorey + plant_raoQ_canopy + plant_raoQ_understorey + CH_m + CH_sd + forest_type + microclim_m_canopy + microclim_var_canopy + PAI_leafoff_canopy + PAI_leafoff_understorey + cover_plant_canopy + cover_plant_understorey |
| biomass_arthropod_1_understorey | year + plant_sr_canopy + plant_sr_understorey + plant_raoQ_canopy + plant_raoQ_understorey + CH_m + CH_sd + forest_type + microclim_m_understorey + microclim_var_understorey + PAI_leafoff_canopy + PAI_leafoff_understorey + cover_plant_canopy + cover_plant_understorey |
| biomass_arthropod_2_canopy | year + plant_sr_canopy + plant_sr_understorey + plant_raoQ_canopy + plant_raoQ_understorey + CH_m + CH_sd + forest_type + microclim_m_canopy + microclim_var_canopy + biomass_arthropod_1_canopy + biomass_arthropod_1_understorey + PAI_leafoff_canopy + PAI_leafoff_understorey + cover_plant_canopy + cover_plant_understorey |
| biomass_arthropod_2_understorey | year + plant_sr_canopy + plant_sr_understorey + plant_raoQ_canopy + plant_raoQ_understorey + CH_m + CH_sd + forest_type + microclim_m_understorey + microclim_var_understorey + biomass_arthropod_1_canopy + biomass_arthropod_1_understorey + PAI_leafoff_canopy + PAI_leafoff_understorey + cover_plant_canopy + cover_plant_understorey |
| biomass_bird_1_total | year + plant_sr_canopy + plant_sr_understorey + plant_raoQ_canopy + plant_raoQ_understorey + CH_m + CH_sd + forest_type + microclim_m_canopy + microclim_var_canopy + microclim_m_understorey + microclim_var_understorey + PAI_leafoff_canopy + PAI_leafoff_understorey + cover_plant_canopy + cover_plant_understorey |
| biomass_bird_2_total | year + plant_sr_canopy + plant_sr_understorey + plant_raoQ_canopy + plant_raoQ_understorey + CH_m + CH_sd + forest_type + microclim_m_canopy + microclim_var_canopy + microclim_m_understorey + microclim_var_understorey + biomass_arthropod_1_canopy + biomass_arthropod_1_understorey + biomass_arthropod_2_canopy + biomass_arthropod_2_understorey + PAI_leafoff_canopy + PAI_leafoff_understorey + cover_plant_canopy + cover_plant_understorey |

| Model | PPP | Response | Predictors | R^2^ |
| --- | --- | --- | --- | --- |
| All arthropods | 0.643 | biomass_arthropod_1_canopy | CH_sd + plant_raoQ_understorey + year | 0.272 |
|  |  | biomass_arthropod_1_understorey | CH_m + microclim_m_understorey + plant_raoQ_understorey + year | 0.152 |
|  |  | biomass_arthropod_2_canopy | biomass_arthropod_1_canopy + PAI_leafoff_understorey + plant_raoQ_understorey + plant_sr_understorey + year | 0.759 |
|  |  | biomass_arthropod_2_understorey | biomass_arthropod_1_understorey + CH_m + microclim_m_understorey + n_plant_canopy + plant_sr_canopy + year | 0.653 |
|  |  | biomass_bird_1_total | plant_raoQ_understorey + year | 0.190 |
|  |  | biomass_bird_2_total | microclim_m_canopy + PAI_leafoff_understorey + year | 0.141 |
|  | 0.532 | n_arthropod_1_canopy | PAI_leafoff_canopy + microclim_m_canopy + plant_raoQ_canopy + year | 0.283 |
|  |  | n_arthropod_1_understorey | microclim_m_understorey + n_plant_canopy + PAI_leafoff_canopy + plant_sr_understorey + year | 0.399 |
|  |  | n_arthropod_2_canopy | CH_m + microclim_m_canopy + n_arthropod_1_canopy + PAI_leafoff_canopy + year | 0.407 |
|  |  | n_arthropod_2_understorey | CH_m + microclim_m_understorey + n_arthropod_1_canopy + n_arthropod_1_understorey + n_plant_canopy + year | 0.538 |
|  |  | n_bird_1_total | microclim_m_canopy + microclim_var_understorey + plant_raoQ_understorey + year | 0.151 |
|  |  | n_bird_2_total | CH_m + CH_sd + microclim_m_canopy + n_arthropod_2_understorey + plant_sr_canopy + year | 0.307 |
| Filtered arthropods | 0.547 | biomass_arthropod_1_canopy | CH_sd + year | 0.256 |
|  |  | biomass_arthropod_1_understorey | CH_m + microclim_m_understorey + PAI_leafoff_understorey + plant_raoQ_understorey + year | 0.176 |
|  |  | biomass_arthropod_2_canopy | biomass_arthropod_1_canopy + PAI_leafoff_understorey + plant_raoQ_understorey + PAI_leafoff_canopy + year | 0.653 |
|  |  | biomass_arthropod_2_understorey | biomass_arthropod_1_understorey + CH_m + microclim_var_understorey + n_plant_canopy + plant_sr_canopy + year | 0.600 |
|  |  | biomass_bird_1_total | plant_raoQ_understorey + year | 0.190 |
|  |  | biomass_bird_2_total | microclim_m_canopy + PAI_leafoff_understorey + year | 0.139 |
|  | 0.453 | n_arthropod_1_canopy | plant_raoQ_canopy + plant_raoQ_understorey + year | 0.242 |
|  |  | n_arthropod_1_understorey | microclim_m_understorey + microclim_var_understorey + n_plant_canopy + PAI_leafoff_understorey + plant_raoQ_canopy + year | 0.404 |
|  |  | n_arthropod_2_canopy | microclim_var_canopy + n_arthropod_1_canopy + PAI_leafoff_canopy + year | 0.440 |
|  |  | n_arthropod_2_understorey | n_arthropod_1_canopy + n_arthropod_1_understorey + plant_sr_canopy + year | 0.620 |
|  |  | n_bird_1_total | microclim_m_canopy + microclim_var_understorey + plant_raoQ_understorey + year | 0.173 |
|  |  | n_bird_2_total | CH_m + CH_sd + microclim_m_canopy + microclim_var_canopy + microclim_var_understorey + year | 0.188 |
| Coleoptera | 0.716 | biomass_arthropod_1_canopy | microclim_m_canopy + PAI_leafoff_canopy + year | 0.150 |
|  |  | biomass_arthropod_1_understorey | CH_m + n_plant_canopy + PAI_leafoff_understorey + plant_raoQ_canopy + plant_raoQ_understorey + microclim_m_understorey + year | 0.493 |
|  |  | biomass_arthropod_2_canopy | biomass_arthropod_1_canopy + biomass_arthropod_1_understorey + PAI_leafoff_understorey + plant_raoQ_understorey + plant_sr_canopy + year | 0.454 |
|  |  | biomass_arthropod_2_understorey | biomass_arthropod_1_understorey + CH_m + CH_sd + microclim_var_understorey + plant_raoQ_understorey + plant_sr_canopy + year | 0.750 |
|  |  | biomass_bird_1_total | plant_raoQ_understorey + year | 0.202 |
|  |  | biomass_bird_2_total | microclim_m_canopy + PAI_leafoff_understorey + plant_sr_canopy + year | 0.148 |
|  | 0.569 | n_arthropod_1_canopy | microclim_m_canopy + PAI_leafoff_canopy + PAI_leafoff_understorey + plant_raoQ_understorey + plant_sr_understorey + year | 0.459 |
|  |  | n_arthropod_1_understorey | CH_m + microclim_var_understorey + n_plant_canopy + PAI_leafoff_understorey + plant_raoQ_canopy + year | 0.252 |
|  |  | n_arthropod_2_canopy | microclim_var_canopy + n_arthropod_1_canopy + year | 0.279 |
|  |  | n_arthropod_2_understorey | n_arthropod_1_canopy + n_arthropod_1_understorey + plant_sr_canopy + year | 0.625 |
|  |  | n_bird_1_total | microclim_m_canopy + microclim_var_understorey + plant_raoQ_understorey + year | 0.162 |
|  |  | n_bird_2_total | CH_m + CH_sd + microclim_m_canopy + microclim_var_canopy + microclim_var_understorey + year | 0.209 |

Table S5. Structure of selected models imputed into the SEMs. Since automated selection was done for each model type (all arthropods, filtered and Coleoptera), they have different explanatory variables. PPP values per model and R^2^ values per regression are also represented.

Table S4. Final model structures for effective biomass (top) and abundance (n, bottom) after model selection, together with the R^2^ value per regression resulting from the BSEMs.

Figure S5. Median effective biomass (mg, top) and abundance (number of individuals, bottom) of the four most abundant arthropod suborders per trophic level and vertical layer (canopy vs. understorey). Colours represent the distinct suborders.

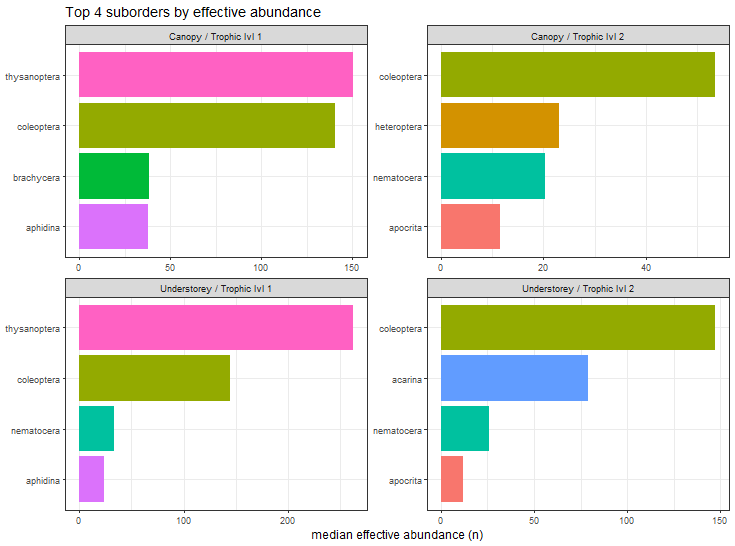

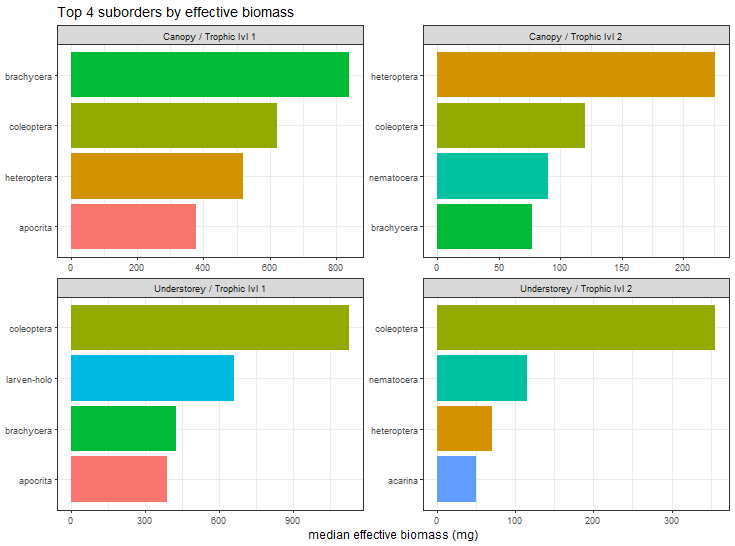

Figure 2. Across all panels, silhouettes represent the concerned taxonomic groups (arthropods and birds), and dark purple hues represent secondary consumers (ladybug and warbler silhouette), while light purple hues represent primary consumers (thrips, weevil (specifically representing herbivore Coleoptera) and wood pigeon). For simplicity, no distinction between biomass and abundance is made in this figure (refer to Fig. 3, S6-S7 for details). Effects are shown with arrows, where positive ones are in light green and a plus (+) sign next to the arrows, negative effects in red and a minus (−) sign, and both negative and positive in light grey and no sign. Panel A represents the generalised findings across biomass and abundance, vertical layers and trophic interactions, with numbers on the top-left corners of the boxes on the left and box line styles, linking to the text boxes on the right, with an explaining summary. This panel refers to the results addressed in the first section of the discussion. Panel B shows the detailed findings covered in the second section of the discussion, from left to right depicted in the same order as addressed in the text. Acronyms: FD (woody plant functional diversity), SR (woody plant species richness), PAI (plant area index), CH (mean plot canopy height), % cover (percentage woody plant cover), CH SD (canopy height standard deviation, i.e. heterogeneity) and symbols: red thermometer and crossed blue water droplet (mean microclimate, i.e. warm-dry axis), red thermometer and blue water droplet with up-down grey arrow (microclimate variability), are used for simplicity.

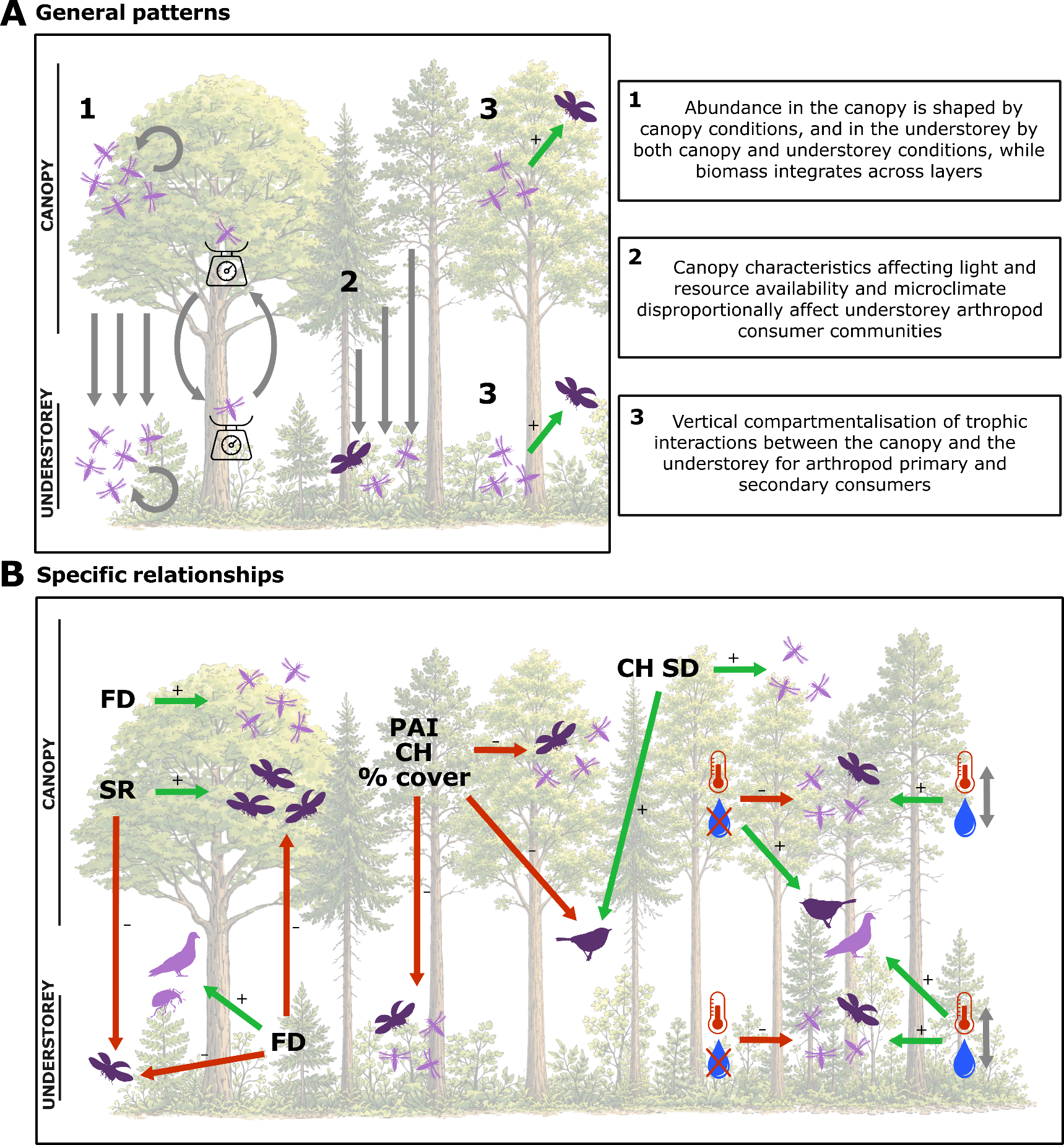

Table S6. Posterior median, 95 and 90 % Equal-Tail credible intervals (upper and lower CrI) for each individual predictor variable for the BSEMs of effective biomass and abundance (n) based on the full arthropod data. Also shown (furthermost right), significance based on whether the 90 % CrI crosses zero (*) or not.

| Parameters | Median | CrI 95 lower | CrI 95 upper | CrI 90 lower | CrI 90 upper |  |
| --- | --- | --- | --- | --- | --- | --- |
| biomass_arthropod_1_canopy~CH_sd | 0.33 | 0.12 | 0.55 | 0.15 | 0.51 | * |
| biomass_arthropod_1_canopy~plant_raoQ_understorey | -0.16 | -0.37 | 0.05 | -0.34 | 0.02 |  |
| biomass_arthropod_1_canopy~year | 0.80 | 0.38 | 1.23 | 0.45 | 1.16 | * |
| biomass_arthropod_1_understorey~CH_m | 0.33 | 0.09 | 0.57 | 0.13 | 0.53 | * |
| biomass_arthropod_1_understorey~microclim_m_understorey | -0.29 | -0.61 | 0.02 | -0.55 | -0.03 | * |
| biomass_arthropod_1_understorey~plant_raoQ_understorey | 0.16 | -0.08 | 0.40 | -0.04 | 0.36 |  |
| biomass_arthropod_1_understorey~year | -0.31 | -1.09 | 0.47 | -0.96 | 0.35 |  |
| biomass_arthropod_2_canopy~biomass_arthropod_1_canopy | 0.68 | 0.55 | 0.81 | 0.57 | 0.79 | * |
| biomass_arthropod_2_canopy~PAI_leafoff_understorey | 0.24 | 0.10 | 0.38 | 0.12 | 0.36 | * |
| biomass_arthropod_2_canopy~plant_raoQ_understorey | -0.21 | -0.34 | -0.09 | -0.32 | -0.11 | * |
| biomass_arthropod_2_canopy~plant_sr_understorey | -0.11 | -0.25 | 0.02 | -0.23 | 0.00 |  |
| biomass_arthropod_2_canopy~year | 0.44 | 0.18 | 0.70 | 0.22 | 0.66 | * |
| biomass_arthropod_2_understorey~biomass_arthropod_1_understorey | 0.31 | 0.15 | 0.47 | 0.18 | 0.44 | * |
| biomass_arthropod_2_understorey~CH_m | -0.43 | -0.59 | -0.26 | -0.56 | -0.29 | * |
| biomass_arthropod_2_understorey~microclim_m_understorey | -0.23 | -0.42 | -0.04 | -0.39 | -0.07 | * |
| biomass_arthropod_2_understorey~n_plant_canopy | -0.58 | -0.75 | -0.41 | -0.72 | -0.44 | * |
| biomass_arthropod_2_understorey~plant_sr_canopy | -0.16 | -0.31 | 0.00 | -0.29 | -0.03 | * |
| biomass_arthropod_2_understorey~year | 0.31 | -0.18 | 0.79 | -0.10 | 0.71 |  |
| biomass_bird_1_total~plant_raoQ_understorey | 0.30 | 0.08 | 0.52 | 0.12 | 0.48 | * |
| biomass_bird_1_total~year | 0.67 | 0.22 | 1.12 | 0.30 | 1.05 | * |
| biomass_bird_2_total~microclim_m_canopy | 0.45 | 0.14 | 0.76 | 0.19 | 0.71 | * |
| biomass_bird_2_total~PAI_leafoff_understorey | 0.26 | 0.02 | 0.50 | 0.06 | 0.45 | * |
| biomass_bird_2_total~year | 0.94 | 0.12 | 1.76 | 0.25 | 1.63 | * |
| n_arthropod_1_canopy~PAI_leafoff_canopy | -0.40 | -0.59 | -0.22 | -0.63 | -0.18 | * |
| n_arthropod_1_canopy~microclim_m_canopy | -0.18 | -0.40 | 0.04 | -0.44 | 0.09 |  |
| n_arthropod_1_canopy~plant_raoQ_canopy | 0.31 | 0.15 | 0.49 | 0.11 | 0.52 | * |
| n_arthropod_1_canopy~year | 0.26 | -0.32 | 0.85 | -0.43 | 0.97 |  |
| n_arthropod_1_understorey~microclim_m_understorey | -0.34 | -0.55 | -0.12 | -0.59 | -0.08 | * |
| n_arthropod_1_understorey~n_plant_canopy | -0.36 | -0.53 | -0.20 | -0.56 | -0.17 | * |
| n_arthropod_1_understorey~PAI_leafoff_canopy | -0.40 | -0.58 | -0.21 | -0.62 | -0.17 | * |
| n_arthropod_1_understorey~plant_sr_understorey | -0.16 | -0.31 | 0.00 | -0.34 | 0.02 | * |
| n_arthropod_1_understorey~year | 0.24 | -0.28 | 0.77 | -0.39 | 0.88 |  |
| n_arthropod_2_canopy~CH_m | 0.06 | -0.11 | 0.23 | -0.14 | 0.27 |  |
| n_arthropod_2_canopy~microclim_m_canopy | -0.26 | -0.51 | -0.03 | -0.56 | 0.02 | * |
| n_arthropod_2_canopy~n_arthropod_1_canopy | 0.42 | 0.09 | 0.74 | 0.02 | 0.81 | * |
| n_arthropod_2_canopy~PAI_leafoff_canopy | 0.16 | -0.04 | 0.35 | -0.08 | 0.39 |  |
| n_arthropod_2_canopy~year | -0.03 | -0.60 | 0.54 | -0.71 | 0.66 |  |
| n_arthropod_2_understorey~CH_m | -0.31 | -0.47 | -0.16 | -0.50 | -0.13 | * |
| n_arthropod_2_understorey~microclim_m_understorey | -0.46 | -0.68 | -0.26 | -0.73 | -0.21 | * |
| n_arthropod_2_understorey~n_arthropod_1_canopy | -0.21 | -0.46 | 0.05 | -0.51 | 0.11 |  |
| n_arthropod_2_understorey~n_arthropod_1_understorey | 0.19 | -0.36 | 0.68 | -0.48 | 0.79 |  |
| n_arthropod_2_understorey~n_plant_canopy | -0.47 | -0.70 | -0.27 | -0.75 | -0.23 | * |
| n_arthropod_2_understorey~year | -0.18 | -0.69 | 0.36 | -0.80 | 0.49 |  |
| n_bird_1_total~microclim_m_canopy | 0.40 | 0.14 | 0.66 | 0.09 | 0.72 | * |
| n_bird_1_total~microclim_var_understorey | 0.18 | 0.04 | 0.32 | 0.01 | 0.34 | * |
| n_bird_1_total~plant_raoQ_understorey | 0.21 | 0.07 | 0.36 | 0.04 | 0.39 | * |
| n_bird_1_total~year | 1.48 | 0.74 | 2.24 | 0.60 | 2.38 | * |
| n_bird_2_total~CH_m | -0.27 | -0.42 | -0.12 | -0.45 | -0.09 | * |
| n_bird_2_total~CH_sd | 0.42 | 0.27 | 0.57 | 0.24 | 0.60 | * |
| n_bird_2_total~microclim_m_canopy | 0.46 | 0.22 | 0.69 | 0.17 | 0.74 | * |
| n_bird_2_total~n_arthropod_2_understorey | 0.30 | 0.14 | 0.46 | 0.11 | 0.49 | * |
| n_bird_2_total~plant_sr_canopy | 0.09 | -0.05 | 0.23 | -0.08 | 0.26 |  |
| n_bird_2_total~year | 0.68 | 0.09 | 1.28 | -0.03 | 1.40 | * |

Table S7. Posterior median, 95 and 90 % Equal-Tail credible intervals (upper and lower CrI) for each individual predictor variable for the BSEMs of effective biomass and abundance (n) based on the filtered arthropod data. Also shown (furthermost right), significance based on whether the 90 % CrI crosses zero (*) or not.

| Parameters | Median | CrI 95 lower | CrI 95 upper | CrI 90 lower | CrI 90 upper |  |
| --- | --- | --- | --- | --- | --- | --- |
| biomass_arthropod_1_canopy~CH_sd | 0.34 | 0.16 | 0.52 | 0.13 | 0.56 | * |
| biomass_arthropod_1_canopy~year | 0.78 | 0.42 | 1.14 | 0.35 | 1.21 | * |
| biomass_arthropod_1_understorey~CH_m | 0.32 | 0.13 | 0.52 | 0.09 | 0.56 | * |
| biomass_arthropod_1_understorey~microclim_m_understorey | -0.31 | -0.57 | -0.05 | -0.62 | 0.01 | * |
| biomass_arthropod_1_understorey~PAI_leafoff_understorey | -0.18 | -0.38 | 0.03 | -0.42 | 0.07 |  |
| biomass_arthropod_1_understorey~plant_raoQ_understorey | 0.24 | 0.03 | 0.44 | -0.01 | 0.48 | * |
| biomass_arthropod_1_understorey~year | -0.47 | -1.12 | 0.17 | -1.25 | 0.30 |  |
| biomass_arthropod_2_canopy~biomass_arthropod_1_canopy | 0.59 | 0.46 | 0.71 | 0.44 | 0.74 | * |
| biomass_arthropod_2_canopy~PAI_leafoff_understorey | 0.19 | 0.06 | 0.32 | 0.04 | 0.34 | * |
| biomass_arthropod_2_canopy~plant_raoQ_understorey | -0.24 | -0.37 | -0.11 | -0.39 | -0.09 | * |
| biomass_arthropod_2_canopy~PAI_leafoff_canopy | -0.15 | -0.28 | -0.02 | -0.30 | 0.00 | * |
| biomass_arthropod_2_canopy~year | 0.48 | 0.22 | 0.73 | 0.17 | 0.78 | * |
| biomass_arthropod_2_understorey~biomass_arthropod_1_understorey | 0.38 | 0.25 | 0.52 | 0.22 | 0.55 | * |
| biomass_arthropod_2_understorey~CH_m | -0.41 | -0.56 | -0.25 | -0.59 | -0.22 | * |
| biomass_arthropod_2_understorey~microclim_var_understorey | 0.11 | -0.02 | 0.25 | -0.05 | 0.28 |  |
| biomass_arthropod_2_understorey~n_plant_canopy | -0.54 | -0.70 | -0.37 | -0.74 | -0.34 | * |
| biomass_arthropod_2_understorey~plant_sr_canopy | -0.20 | -0.36 | -0.05 | -0.39 | -0.02 | * |
| biomass_arthropod_2_understorey~year | 0.81 | 0.47 | 1.14 | 0.40 | 1.20 | * |
| biomass_bird_1_total~plant_raoQ_understorey | 0.30 | 0.11 | 0.49 | 0.08 | 0.52 | * |
| biomass_bird_1_total~year | 0.67 | 0.30 | 1.05 | 0.23 | 1.13 | * |
| biomass_bird_2_total~microclim_m_canopy | 0.44 | 0.18 | 0.70 | 0.13 | 0.75 | * |
| biomass_bird_2_total~PAI_leafoff_understorey | 0.26 | 0.06 | 0.46 | 0.02 | 0.50 | * |
| biomass_bird_2_total~year | 0.93 | 0.23 | 1.62 | 0.10 | 1.75 | * |
| n_arthropod_1_canopy~plant_raoQ_canopy | 0.44 | 0.25 | 0.63 | 0.21 | 0.66 | * |
| n_arthropod_1_canopy~plant_raoQ_understorey | -0.23 | -0.39 | -0.07 | -0.42 | -0.04 | * |
| n_arthropod_1_canopy~year | 0.54 | 0.18 | 0.90 | 0.10 | 0.97 | * |
| n_arthropod_1_understorey~microclim_m_understorey | -0.23 | -0.42 | -0.04 | -0.45 | 0.00 | * |
| n_arthropod_1_understorey~microclim_var_understorey | 0.26 | 0.11 | 0.40 | 0.09 | 0.43 | * |
| n_arthropod_1_understorey~n_plant_canopy | -0.08 | -0.24 | 0.07 | -0.27 | 0.10 |  |
| n_arthropod_1_understorey~PAI_leafoff_understorey | -0.29 | -0.46 | -0.13 | -0.49 | -0.09 | * |
| n_arthropod_1_understorey~plant_raoQ_canopy | 0.39 | 0.20 | 0.57 | 0.17 | 0.60 | * |
| n_arthropod_1_understorey~year | 0.73 | 0.21 | 1.25 | 0.11 | 1.35 | * |
| n_arthropod_2_canopy~microclim_var_canopy | 0.31 | 0.13 | 0.48 | 0.10 | 0.52 | * |
| n_arthropod_2_canopy~n_arthropod_1_canopy | 0.47 | 0.24 | 0.71 | 0.19 | 0.75 | * |
| n_arthropod_2_canopy~PAI_leafoff_canopy | -0.17 | -0.34 | 0.00 | -0.37 | 0.03 | * |
| n_arthropod_2_canopy~year | 1.02 | 0.59 | 1.46 | 0.50 | 1.55 | * |
| n_arthropod_2_understorey~n_arthropod_1_canopy | -0.29 | -0.52 | -0.05 | -0.58 | 0.00 | * |
| n_arthropod_2_understorey~n_arthropod_1_understorey | 0.93 | 0.60 | 1.26 | 0.53 | 1.33 | * |
| n_arthropod_2_understorey~plant_sr_canopy | -0.31 | -0.44 | -0.18 | -0.47 | -0.15 | * |
| n_arthropod_2_understorey~year | -0.07 | -0.40 | 0.26 | -0.46 | 0.32 |  |
| n_bird_1_total~microclim_m_canopy | 0.45 | 0.19 | 0.71 | 0.13 | 0.76 | * |
| n_bird_1_total~microclim_var_understorey | 0.23 | 0.04 | 0.41 | 0.00 | 0.45 | * |
| n_bird_1_total~plant_raoQ_understorey | 0.22 | 0.08 | 0.37 | 0.05 | 0.40 | * |
| n_bird_1_total~year | 1.66 | 0.86 | 2.45 | 0.70 | 2.60 | * |
| n_bird_2_total~CH_m | -0.27 | -0.44 | -0.10 | -0.47 | -0.07 | * |
| n_bird_2_total~CH_sd | 0.44 | 0.28 | 0.60 | 0.25 | 0.63 | * |
| n_bird_2_total~microclim_m_canopy | 0.36 | 0.10 | 0.62 | 0.05 | 0.67 | * |
| n_bird_2_total~microclim_var_canopy | -0.08 | -0.24 | 0.08 | -0.27 | 0.11 |  |
| n_bird_2_total~microclim_var_understorey | 0.07 | -0.11 | 0.26 | -0.15 | 0.29 |  |
| n_bird_2_total~year | 0.70 | -0.12 | 1.51 | -0.27 | 1.68 |  |

| Parameters | Median | CrI 95 lower | CrI 95 upper | CrI 90 lower | CrI 90 upper |  |
| --- | --- | --- | --- | --- | --- | --- |
| biomass_arthropod_1_canopy~microclim_m_canopy | -0.33 | -0.56 | -0.10 | -0.61 | -0.05 | * |
| biomass_arthropod_1_canopy~PAI_leafoff_canopy | -0.17 | -0.35 | 0.01 | -0.38 | 0.04 |  |
| biomass_arthropod_1_canopy~year | -1.28 | -1.91 | -0.65 | -2.04 | -0.53 | * |
| biomass_arthropod_1_understorey~CH_m | -0.17 | -0.33 | -0.01 | -0.36 | 0.02 | * |
| biomass_arthropod_1_understorey~n_plant_canopy | -0.55 | -0.70 | -0.40 | -0.73 | -0.37 | * |
| biomass_arthropod_1_understorey~PAI_leafoff_understorey | -0.36 | -0.50 | -0.21 | -0.53 | -0.19 | * |
| biomass_arthropod_1_understorey~plant_raoQ_canopy | 0.11 | -0.04 | 0.26 | -0.07 | 0.29 |  |
| biomass_arthropod_1_understorey~plant_raoQ_understorey | 0.35 | 0.20 | 0.50 | 0.17 | 0.53 | * |
| biomass_arthropod_1_understorey~microclim_m_understorey | -0.13 | -0.32 | 0.06 | -0.36 | 0.10 |  |
| biomass_arthropod_1_understorey~year | -0.10 | -0.59 | 0.39 | -0.68 | 0.49 |  |
| biomass_arthropod_2_canopy~biomass_arthropod_1_canopy | 0.42 | 0.24 | 0.61 | 0.20 | 0.65 | * |
| biomass_arthropod_2_canopy~biomass_arthropod_1_understorey | 0.29 | 0.09 | 0.50 | 0.05 | 0.54 | * |
| biomass_arthropod_2_canopy~PAI_leafoff_understorey | 0.14 | -0.07 | 0.35 | -0.11 | 0.40 |  |
| biomass_arthropod_2_canopy~plant_raoQ_understorey | -0.27 | -0.45 | -0.08 | -0.49 | -0.04 | * |
| biomass_arthropod_2_canopy~plant_sr_canopy | 0.32 | 0.13 | 0.52 | 0.09 | 0.56 | * |
| biomass_arthropod_2_canopy~year | -0.24 | -0.58 | 0.10 | -0.65 | 0.17 |  |
| biomass_arthropod_2_understorey~biomass_arthropod_1_understorey | 0.68 | 0.55 | 0.80 | 0.53 | 0.83 | * |
| biomass_arthropod_2_understorey~CH_m | -0.19 | -0.31 | -0.07 | -0.33 | -0.05 | * |
| biomass_arthropod_2_understorey~CH_sd | 0.25 | 0.12 | 0.38 | 0.09 | 0.40 | * |
| biomass_arthropod_2_understorey~microclim_var_understorey | 0.13 | 0.03 | 0.23 | 0.01 | 0.25 | * |
| biomass_arthropod_2_understorey~plant_raoQ_understorey | -0.15 | -0.27 | -0.04 | -0.29 | -0.02 | * |
| biomass_arthropod_2_understorey~plant_sr_canopy | -0.21 | -0.33 | -0.09 | -0.35 | -0.07 | * |
| biomass_arthropod_2_understorey~year | 0.29 | 0.04 | 0.55 | -0.01 | 0.60 | * |
| biomass_bird_1_total~plant_raoQ_understorey | 0.33 | 0.14 | 0.51 | 0.10 | 0.55 | * |
| biomass_bird_1_total~year | 0.67 | 0.29 | 1.06 | 0.22 | 1.13 | * |
| biomass_bird_2_total~microclim_m_canopy | 0.44 | 0.18 | 0.70 | 0.13 | 0.75 | * |
| biomass_bird_2_total~PAI_leafoff_understorey | 0.18 | -0.07 | 0.42 | -0.12 | 0.47 |  |
| biomass_bird_2_total~plant_sr_canopy | 0.14 | -0.11 | 0.39 | -0.16 | 0.43 |  |
| biomass_bird_2_total~year | 0.92 | 0.23 | 1.60 | 0.10 | 1.74 | * |
| n_arthropod_1_canopy~microclim_m_canopy | -0.21 | -0.40 | -0.02 | -0.44 | 0.01 | * |
| n_arthropod_1_canopy~PAI_leafoff_canopy | -0.37 | -0.53 | -0.21 | -0.56 | -0.17 | * |
| n_arthropod_1_canopy~PAI_leafoff_understorey | 0.35 | 0.16 | 0.53 | 0.13 | 0.57 | * |
| n_arthropod_1_canopy~plant_raoQ_understorey | -0.31 | -0.45 | -0.16 | -0.48 | -0.13 | * |
| n_arthropod_1_canopy~plant_sr_understorey | -0.10 | -0.26 | 0.06 | -0.29 | 0.09 |  |
| n_arthropod_1_canopy~year | -1.05 | -1.55 | -0.55 | -1.65 | -0.45 | * |
| n_arthropod_1_understorey~CH_m | -0.32 | -0.50 | -0.14 | -0.55 | -0.10 | * |
| n_arthropod_1_understorey~microclim_var_understorey | 0.17 | -0.01 | 0.35 | -0.04 | 0.39 |  |
| n_arthropod_1_understorey~n_plant_canopy | -0.26 | -0.46 | -0.07 | -0.50 | -0.03 | * |
| n_arthropod_1_understorey~PAI_leafoff_understorey | -0.30 | -0.50 | -0.10 | -0.54 | -0.05 | * |
| n_arthropod_1_understorey~plant_raoQ_canopy | 0.30 | 0.12 | 0.48 | 0.08 | 0.52 | * |
| n_arthropod_1_understorey~year | 0.44 | -0.01 | 0.88 | -0.09 | 0.97 |  |
| n_arthropod_2_canopy~microclim_var_canopy | 0.32 | 0.15 | 0.49 | 0.11 | 0.53 | * |
| n_arthropod_2_canopy~n_arthropod_1_canopy | 0.48 | 0.24 | 0.72 | 0.19 | 0.77 | * |
| n_arthropod_2_canopy~year | 0.82 | 0.33 | 1.30 | 0.24 | 1.40 | * |
| n_arthropod_2_understorey~n_arthropod_1_canopy | -0.12 | -0.26 | 0.03 | -0.29 | 0.06 |  |
| n_arthropod_2_understorey~n_arthropod_1_understorey | 0.67 | 0.49 | 0.85 | 0.45 | 0.89 | * |
| n_arthropod_2_understorey~plant_sr_canopy | -0.21 | -0.33 | -0.09 | -0.36 | -0.06 | * |
| n_arthropod_2_understorey~year | 0.15 | -0.12 | 0.42 | -0.17 | 0.47 |  |
| n_bird_1_total~microclim_m_canopy | 0.43 | 0.17 | 0.70 | 0.11 | 0.75 | * |
| n_bird_1_total~microclim_var_understorey | 0.22 | 0.04 | 0.41 | 0.00 | 0.45 | * |
| n_bird_1_total~plant_raoQ_understorey | 0.21 | 0.06 | 0.35 | 0.03 | 0.38 | * |
| n_bird_1_total~year | 1.62 | 0.82 | 2.42 | 0.67 | 2.57 | * |
| n_bird_2_total~CH_m | -0.31 | -0.47 | -0.15 | -0.50 | -0.12 | * |
| n_bird_2_total~CH_sd | 0.47 | 0.31 | 0.62 | 0.28 | 0.66 | * |
| n_bird_2_total~microclim_m_canopy | 0.38 | 0.13 | 0.64 | 0.07 | 0.69 | * |
| n_bird_2_total~microclim_var_canopy | -0.07 | -0.22 | 0.09 | -0.25 | 0.12 |  |

Table S8. Posterior median, 95 and 90 % Equal-Tail credible intervals (upper and lower CrI) for each individual predictor variable for the BSEMs of effective biomass and abundance (n) based on the Coleoptera data. Also shown (furthermost right), significance based on whether the 90 % CrI crosses zero (*) or not.

Figure 3. Equal-Tail credible intervals (CrIs) at 90 % (thick lines) and 95 % (thin lines, extending from thick lines), and median (central dot), for the coefficients of the variables in the BSEM. The top six panels show the predictor variable CrIs for the full biomass model and the bottom six panels those for the full abundance model. The title for each panel represents the specific regression’s predictor variable (ie. all arthropod biomass belonging to trophic level one on the top left). Coloured lines represent variables measured in the canopy (light green), understorey (dark green), or at the plot level (blue), while grey lines represent CrIs crossing zero at 90 % CrI (non-significant).

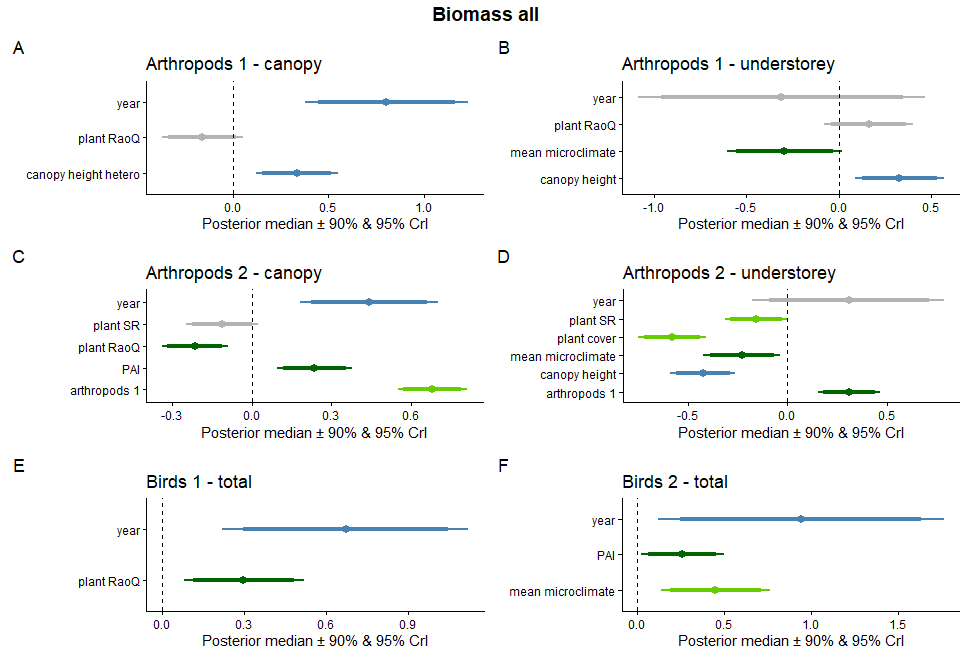

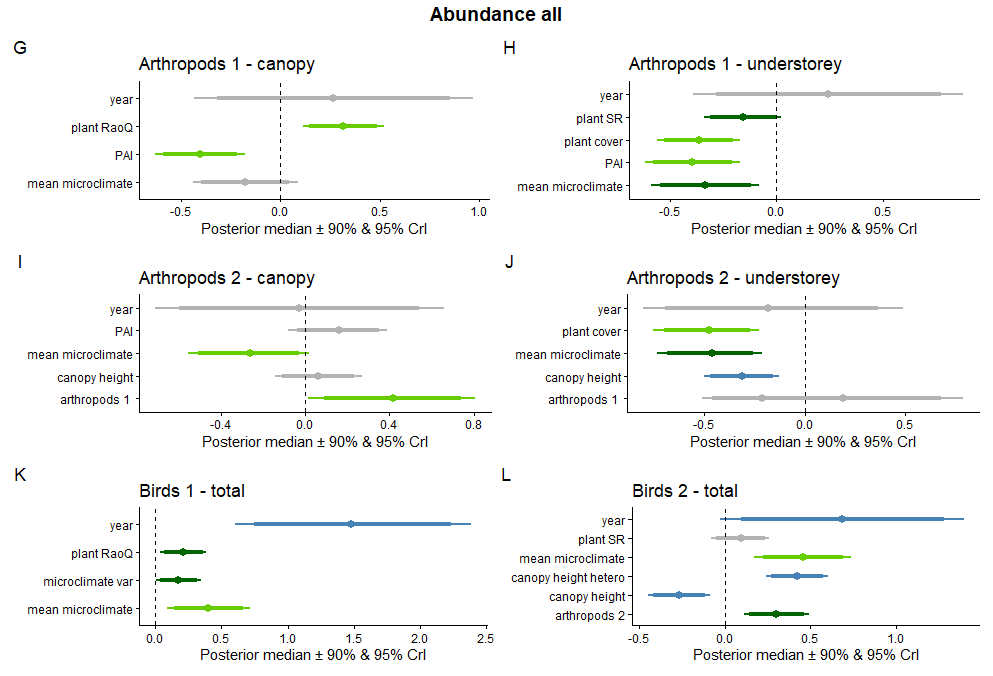

| n_bird_2_total~microclim_var_understorey | 0.06 | -0.12 | 0.24 | -0.16 | 0.27 |
| --- | --- | --- | --- | --- | --- |
| n_bird_2_total~year | 0.73 | -0.07 | 1.54 | -0.24 | 1.69 |

Figure S6. Equal-Tail credible intervals (CrIs) at 90 % (thick lines) and 95 % (thin lines, extending from thick lines), and median (central dot), for the coefficients of the variables in the BSEM. The top six panels show the predictor variable CrIs for the filtered biomass model and the bottom six panels those for the filtered abundance model. The title for each panel represents the specific regression’s predictor variable (ie. filtered arthropod biomass belonging to trophic level one on the top left). Coloured lines represent variables measured in the canopy (light green), understorey (dark green), or at the plot level (blue), while grey lines represent CrIs crossing zero at 90 % CrI (non-significant).

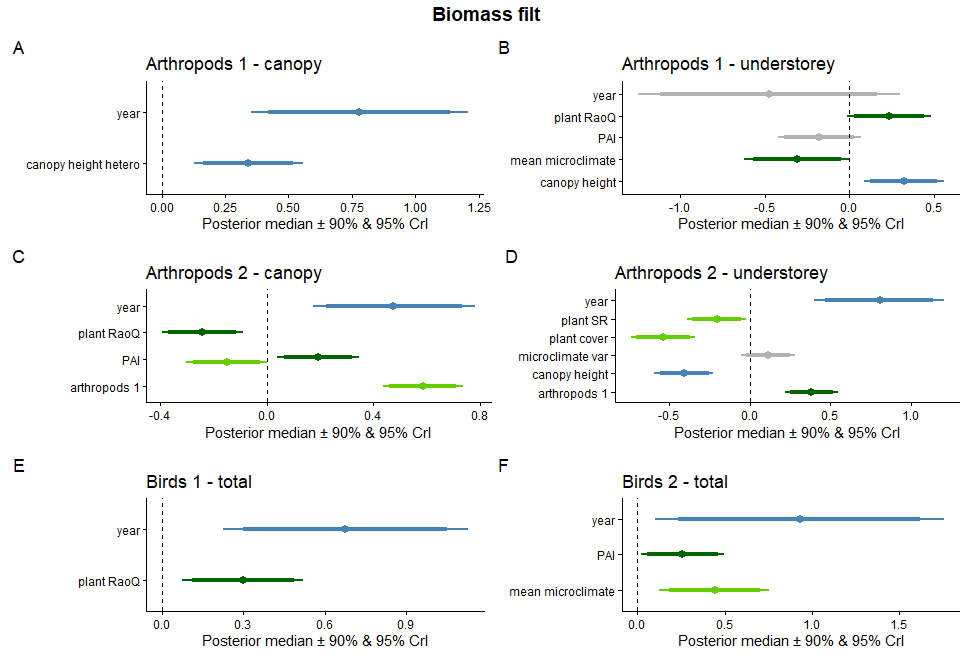

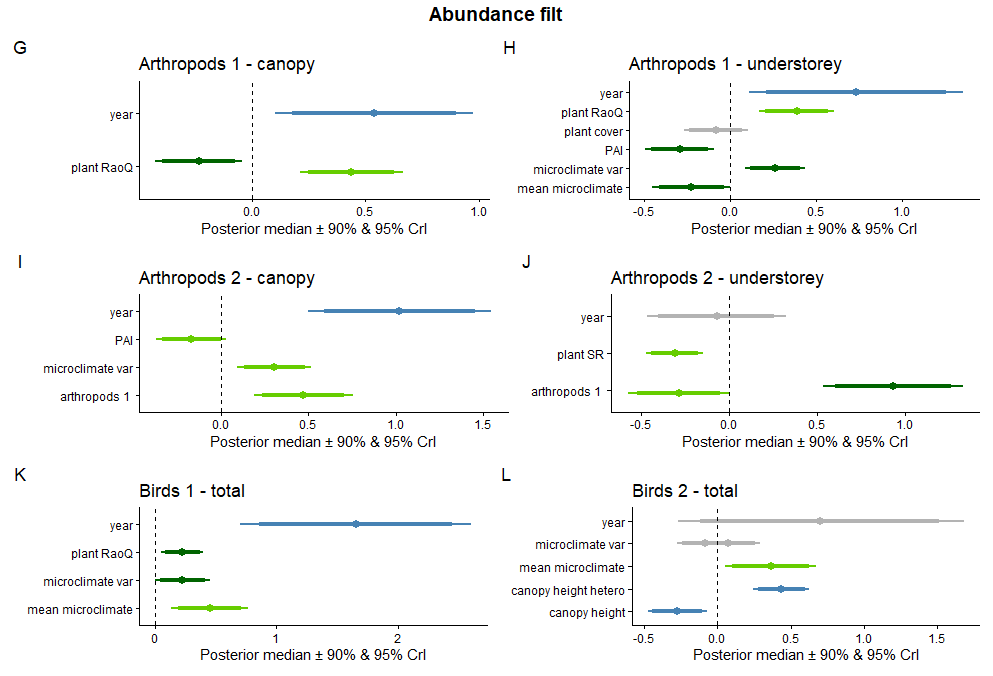

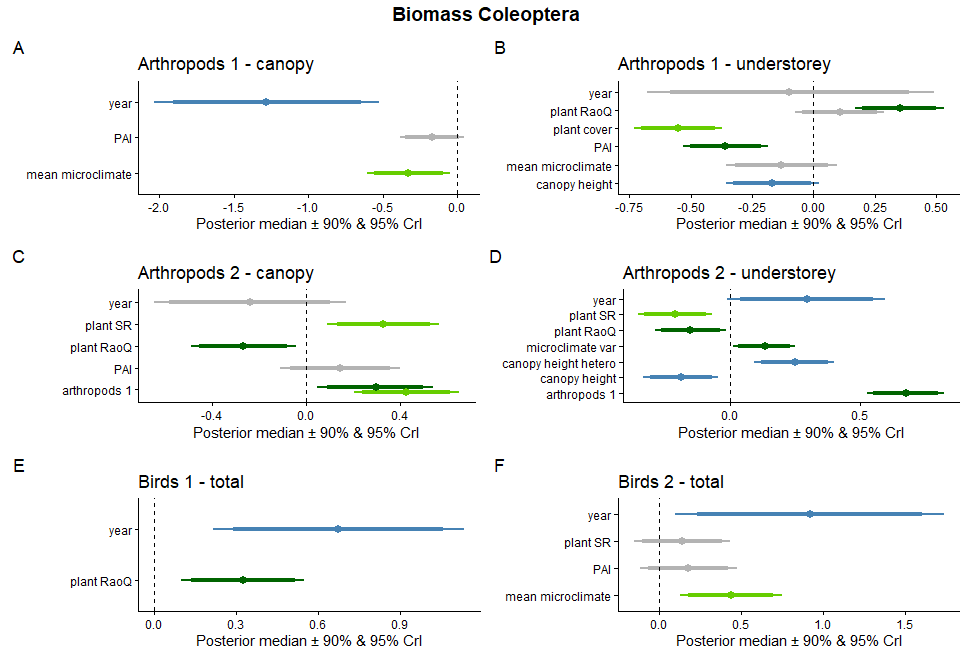

Figure S7. Equal-Tail credible intervals (CrIs) at 90 % (thick lines) and 95 % (thin lines, extending from thick lines), and median (central dot), for the coefficients of the variables in the BSEM. The top six panels show the predictor variable CrIs for the Coleoptera biomass model and the bottom six panels those for the Coleoptera abundance model. The title for each panel represents the specific regression’s predictor variable (ie. Coleoptera biomass belonging to trophic level one on the top left). Coloured lines represent variables measured in the canopy (light green), understorey (dark green), or at the plot level (blue), while grey lines represent CrIs crossing zero at 90 % CrI (non-significant).

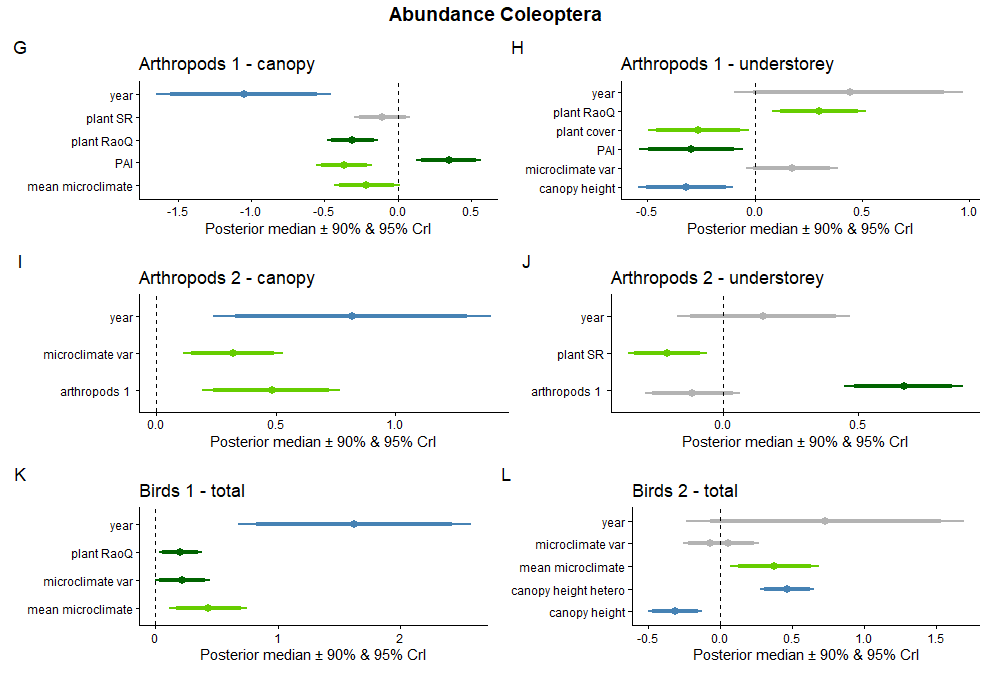
